# In *Campylobacter jejuni* a new type of chaperone receives heme *b* from ferrochelatase

**DOI:** 10.1101/2023.03.30.534706

**Authors:** Jordi Zamarreño Beas, Marco A.M. Videira, Val Karavaeva, Frederico S. Lourenço, Mafalda R. Almeida, Filipa Sousa, Lígia M. Saraiva

**Affiliations:** Instituto de Tecnologia Química e Biológica António Xavier, Universidade Nova de Lisboa, 2780-157 Oeiras, Portugal; Department of Functional and Evolutionary Ecology, University of Vienna, Djerassiplatz 1, 1030 Wien, Austria

## Abstract

Intracellular heme formation and trafficking are fundamental processes in living organisms. Three biogenesis pathways are used by bacteria and archaea to produce iron protoporphyrin IX (heme *b*) that diverge after the formation of the common intermediate uroporphyrinogen III (uro’gen III). In this work, we identify and provide a detailed characterization of the enzymes involved in the transformation of uro’gen III into heme. We show that in this organism operates the protoporphyrin-dependent pathway (PPD pathway), in which the last reaction is the incorporation of ferrous iron into the porphyrin ring by the ferrochelatase enzyme. In general, following this final reaction, little is known about how the formed heme *b* reaches the target proteins. In particular, the chaperons that are thought to be required to traffic heme for incorporation into hemeproteins to avoid the cytotoxicity associated to free heme, remain largely unidentified. We identified in *C. jejuni* a chaperon-like protein, named CgdH2, that binds heme with a dissociation constant of 4.9 ± 1.0 µM, a binding that is impaired upon mutation of residues histidine 45 and 133. We show that *C. jejuni* CgdH2 establishes protein-protein interactions with ferrochelatase, which should enable for the observed transfer of heme from ferrochelatase to CgdH2. Phylogenetic analysis revealed that *C. jejuni* CgdH2 is evolutionarily distinct from the currently known chaperones. Therefore, CgdH2 is a novel chaperone and the first protein identified as an acceptor of the intracellularly formed heme, thus enlarging our understanding of bacterial heme homeostasis.

## Introduction

*Campylobacter jejuni* is a Gram-negative microaerophilic Epsilon proteobacterium responsible for a high percentage of diarrheal infections worldwide, usually caused by ingestion of poultry meat contaminated with strains whose resistance to conventional antibiotics has been increasing (Jadeja and Worrich 2022).

Heme is an iron-porphyrin cofactor essential to several life associated cellular processes such as, respiration, signaling, RNA processing, redox catalysis, stress responses and cellular differentiation. Like almost all other living systems, bacteria require heme for regular functioning and usually express systems dedicated to synthesizing and/or acquiring it from the host.

Most prokaryotes are able to endogenously produce heme *b* (aka heme) through a multi-step process named the heme biosynthesis pathway (Dailey et al. 2017). Three pathways are so known to occur in bacteria and archaea: the protoporphyrin dependent pathway (PPD), the coproporphyrin dependent pathway (CPD), and the siroheme dependent pathway (SHD). All pathways share as the first common precursor the 5-aminolevulinic acid (ALA) molecule, that is converted to uroporphyrinogen III (uro’gen III) through the action of three enzymes: PbgS, HmbS and UroS (Heinemann, Jahn, and Jahn 2008). The following conversion of uro’gen III to heme occurs according to one of three pathways depending on the organism, which are described in detail in several reviews (e.g., Dailey and Medlock 2022; Zamarreño Beas, Videira, and Saraiva 2022).

Most Gram-negative bacteria utilize the PPD pathway, formerly known as the classic pathway, which was the first identified heme biosynthesis route. The PPD pathway starts by the decarboxylation of the acetic acid groups of uro’gen III forming coproporphyrinogen III, followed by decarboxylation of two of the propionic acid chains of coproporphyrinogen III to produce protoporphyrinogen IX. The first decarboxylation reaction is performed by the enzyme uro’gen III decarboxylase (UroD). The second reaction is done by coproporphyrinogen decarboxylase (CgdC) that requires molecular oxygen, or by the oxygen-independent coproporphyrinogen dehydrogenase CgdH (previously named HemN or HemZ). Besides the *bona fide* CgdHs that possess coproporphyrinogen III dehydrogenase activity (Layer et al. 2003), bacteria may contain other *cgdH-*like genes with different functions (Cheng et al. 2022). This is the case of the HemW-like proteins, which are chaperons involved in heme transfer, or have enzyme activities such as methyltransferases and cyclopropanases (Cheng et al. 2022). The protoporphyrinogen IX formed by CgdH is converted to protoporphyrin IX by one of the membrane-associated enzymes PgoX, PgdH1 and PgdH2. PgoX is an oxygen-dependent proto’gen oxidase that contains a tightly non-covalently bound FAD, which was studied in *Acinetobacter* and *Synechocystis* spp. (Dailey et al. 2017; Kobayashi et al. 2014). PgdH1 and PgdH2, that share low amino sequence similarity, are considered protoporphyrinogen dehydrogenases as they require anoxic conditions to produce protoporphyrin IX. PgdH1 is a FMN containing protein that interacts with the cellular respiratory chain. Although PgdH2 is the less-characterized enzyme and it is not known whether it requires specific cofactors, it is the most common protoporphyrinogen dehydrogenase in Gram-negative heme synthesizing bacteria (> 60% occurrence). Finally, in the last step of the heme pathway, iron is inserted into protoporphyrin IX to produce protoheme IX (heme *b*) via the enzyme protoporphyrin ferrochelatase (PpfC).

*C. jejuni* also contains a heme uptake system that includes a TonB-dependent heme receptor (ChuA), a heme ABC transporter permease (ChuB), a heme ABC transporter ATP-binding protein (ChuC), a periplasmic heme-binding protein (ChuD), and the heme oxygenase ChuZ encoded by a gene that is divergently transcribed from the *chuABCD* operon. This heme import system contributes to the virulence of *C. jejuni* (Johnson, Gaddy, and DiRita 2016; Richard, Kelley, and Johnson 2019; Ridley et al. 2006). While the heme uptake in *C. jejuni* is well studied, the type of heme biosynthesis pathway that is active is this bacterium remains to be defined. Here, we have identified how *C. jejuni* produces heme intracellularly through a throughout characterization of the enzymes forming the pathway that goes from uro’gen III to heme.

Although the intracellular heme trafficking is a fundamental biological process that needs to be highly controlled to avoid intracellular heme toxicity, its transport and delivery remain largely unknown. In this work, we found a novel heme binding protein that can receive heme from the last enzyme of the PPD pathway, i.e., from ferrochelatase.

## Results

The heme biosynthesis pathways known to date have uroporphyrinogen III (uro’gen III) as the common precursor that is converted to heme *b* through three possible pathways that vary according to the organism (Figure 1). In this work, we sought to identify which pathway *C. jejuni* uses to synthesize heme from uro’gen III. We search the *C. jejuni* NCTC 11168 genome to identify genes encoding proteins putatively involved in the conversion of uro’gen III to heme *b* and found proteins that share significant amino acid sequence similarity with bacterial heme-biosynthesis related proteins. The *C. jejuni* genes were cloned, the recombinant proteins produced, biochemically characterized, and their structures were modelled using AlphaFold2 (Mirdita et al. 2022). The porphyrin products of the enzymatic assays were detected by UV-visible spectroscopy and high-performance liquid chromatography-mass spectrometry (HPLC-MS).

**Figure 1.**
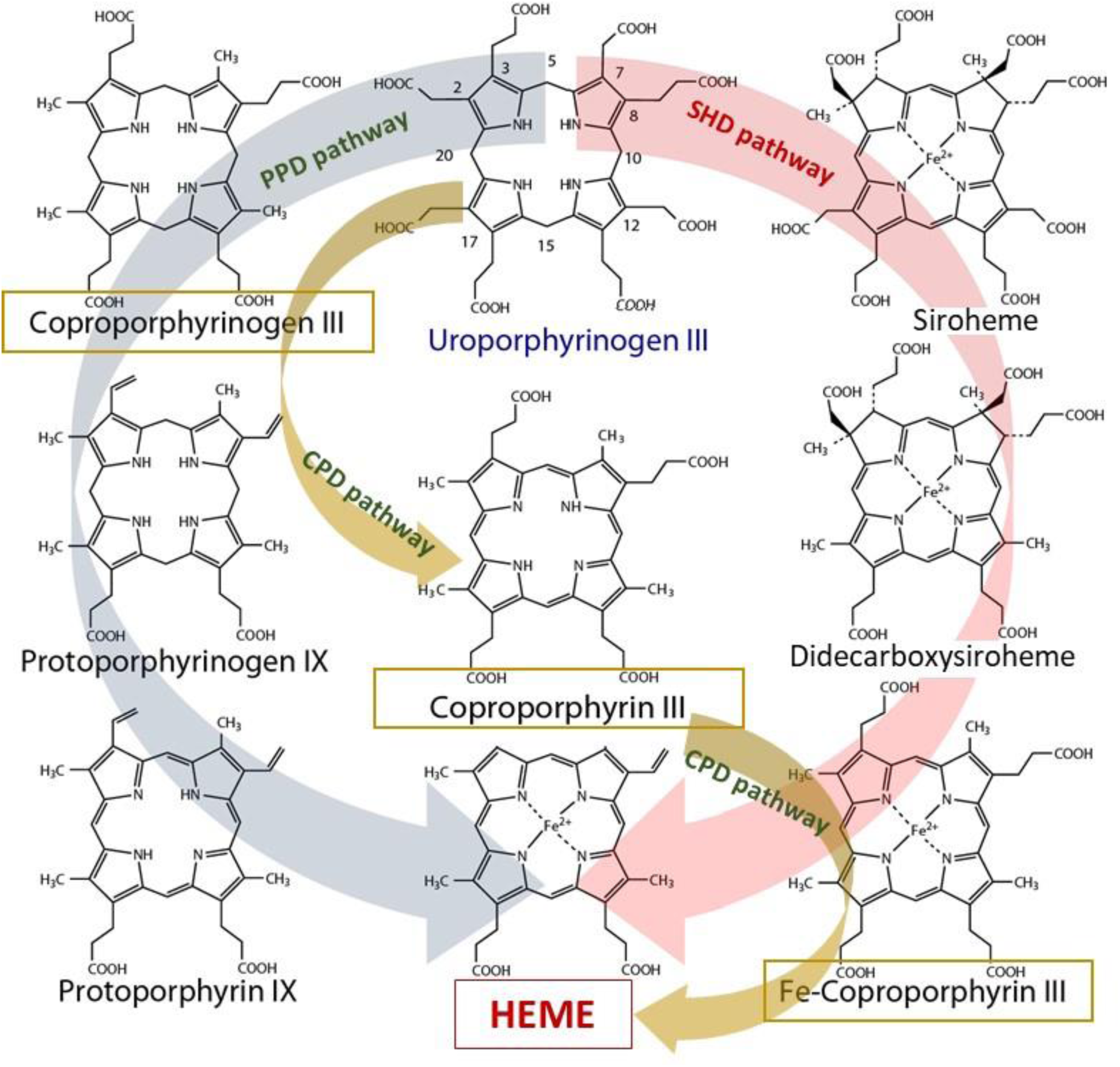
Heme biosynthesis pathways in prokaryotes. Protoporphyrin-dependent (PPD) pathway; coproporphyrin-dependent (CPD) pathway; and siroheme-dependent (SHD) pathway. All routes use four enzymatic steps to convert uroporphyrinogen III to the final product heme. PPD pathway: 1) decarboxylation of the four acetate side chains of uroporphyrinogen III to methyl groups forming coproporphyrinogen III; 2) oxidative decarboxylation of the propionate substituents on the pyrrole rings A and B of coproporphyrinogen III generating the corresponding vinyl groups in protoporphyrinogen IX; 3) oxidation of protoporphyrinogen IX to protoporphyrin IX; and 4) insertion of ferrous iron into protoporphyrin IX for heme formation. CPD pathway: 1) decarboxylation of uroporphyrinogen III to coproporphyrinogen III (identical to the first reaction of the PPD pathway); 2) oxidation of coproporphyrinogen III to coproporphyrin III; 3) insertion of ferrous iron into coproporphyrin III macrocycle forming Fe-coproporphyrin III; and 4) oxidative decarboxylation of the propionate side chains in pyrrole rings A and B of Fe-coproporphyrin III to yield the corresponding vinyl groups of heme. SHD pathway: 1) methylation of uroporphyrinogen III at carbon positions 2 and 7 and insertion of ferrous iron; 2) siroheme methylation at carbon positions 12 and 17 forming didecarboxysiroheme; 3) decarboxylation of didecarboxysiroheme forming Fe-coproporphyrin III; and 4) oxidative removal of two acetate substituents yielding Fe-coproporphyrin III and formation of the vinyl groups of heme.

### *C. jejuni* heme pathway: from uro’gen III to copro’gen III

The first conversion of uro’gen III involves the decarboxylation of the acetic acid groups of uro’gen III promoted by UroD, yielding coproporphyrinogen III (copro’gen III). In *C. jejuni* genome, the *cj1243* gene encodes a protein that shares amino acid sequence identity of ∼45% and ∼38% with UroD of *E. coli* and *B. subtilis*, respectively (Figure 2A).

**Figure 2.**
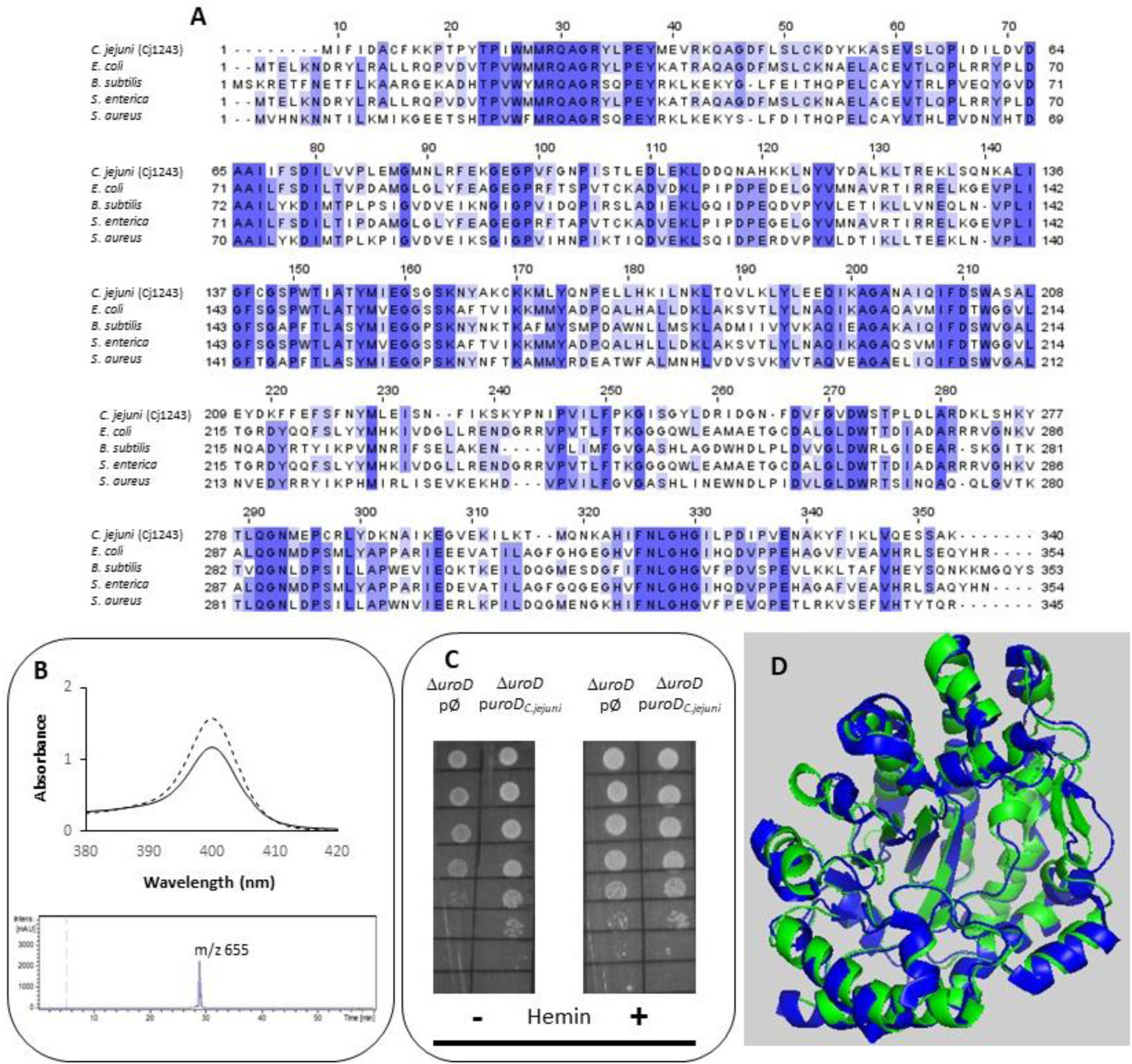
*C. jejuni* UroD is a functional uro’gen III decarboxylase. **A.** Amino acid sequence alignment of *C. jejuni* UroD (Cj1243, UniProt (UP): A8FMU8) and UroD from *E. coli* (UP: P29680), *B. subtilis* (UP: P32395), *S. enterica* (NCBI-Protein ID: AGX12552), and *S. aureus* (UP: A6QI15), using ClustalOmega and edited with Jalview 2.11.2.5 (Waterhouse et al. 2009). Dark blue boxes highlight residues with a percentage identity > 80%. The lighter the blue boxes the lower the percentage identity of the highlighted residues (from 60-0%). **B.** Top-image: UV-visible spectra of the coproporphyrin III standard (dashed line) and the oxidized form of the reaction product of *C. jejuni* UroD, i.e., coproporphyrin III (filled line). Lower image depicts the HPLC-MS extracted-ion chromatogram of the reaction product of the *C. jejuni* UroD reaction. A single peak was observed at the expected m/z of 655 for coproporphyrin III, which corresponds to the oxidized form of coproporphyrinogen III. **C.** *E. coli* Δ*uroD* strain expressing the empty plasmid (pØ) or a plasmid harboring a functional copy of the *C. jejuni* uroD gene (p*uroD_C.jejuni_*), spotted on LB-agar medium in the presence and in the absence of hemin. **D.** Structure model of *C. jejuni* UroD (green), obtained by Alphafold2, superimposed with *B. subtilis* UroD structure (blue, PDB number 2INF), with a RMSD. between the Cα carbons of 1.1.

The Cj1243 protein was produced recombinantly, purified and tested for conversion of uro’gen III to copro’gen III. We obtained the uro’gen III substrate from porphobilinogen using *Desulfovibrio vulgaris* PbgS and UroS enzymes, as described in Materials and Methods. The products of the reaction mixture containing *C. jejuni* Cj1243 and uro’gen III were analyzed by UV-visible spectroscopy and HPLC-MS (Figure 2B). The two typical features of the oxidized form of coproporphyrin III (copro’gen III), namely the development of an absorbance at 400 nm in the visible spectrum and the presence of a mass peak at 655 m/z in the MS, confirmed the formation of copro’gen III.

We determined the *in vivo* activity of *C. jejuni* Cj1243 in a complementation assay using the *E. coli* Δ*uroD* mutant strain transformed with a plasmid that expresses *C. jejuni UroD*. The lack of the *uroD* gene in the *E. coli* Δ*uroD* strain prevents heme synthesis so that the strain only grows in heme-supplemented medium (Săsărman et al. 1975). Figure 2C and Supplementary Figure S1 show that the expression of *C. jejuni* Cj1243 in *E. coli* Δ*uroD*, in the absence of any external source of heme, suppressed the growth defect of *E. coli* Δ*uroD,* i.e., it restored the heme biosynthesis capacity of the mutant strain.

The predicted structure of Cj1243 obtained using AlphaFold2 (Figure 2D) displays a high degree of confidence, and the protein shares significant structural similarity with bacterial homologs. In particular, Cj1243 predicted structure aligns with that of *B. subtilis* UroD with a Root-Mean-Square Distance (RMSD) of 1.1 and retains the main structural features of the *Bacillus* protein. Taken together these results prove that Cj1243 is an uro’gen decarboxylase UroD active in *C. jejuni*.

### *C. jejuni* heme pathway: from copro’gen III to proto’gen IX

Conversion of coprógen III to protoporphyrinogen IX (protógen IX) requires the oxidative decarboxylation of the propionate vinyl groups of rings A and B of coprógen III to yield the vinyl groups of protógen IX. In bacteria, this reaction can be done by two distinct enzymes, namely the aerobic manganese-containing copro’gen III decarboxylase (CgdC) or the anaerobic coproporphyrinogen dehydrogenase CgdH (Cheng et al. 2022; Dailey et al. 2017). BLAST searches of the *C. jejuni* genome did not identify genes encoding bacterial homologues of CgdC but showed that it contains three CgdH-like proteins encoded by *cj0992c*, *cj0363c* and *cj0580c*.

*C. jejuni* Cj0992c and Cj0363c have a similar number of amino acid residues, namely 451 and 448 amino acid residues, respectively, while Cj0580c is much shorter with only 355 amino acid residues. *C. jejuni* Cj0992 shares the highest similarity with *E. coli* CgdH (45% identity, 63% similarity, 98% coverage). *C. jejuni* Cj0363c and Cj0580c share lower similarity with *E. coli* CgdH (26-29% identity, 41-48% similarity, ∼40% coverage) (Figure 3A). The *E. coli* CgdH motifs for the coproporphyrinogen dehydrogenase activity (Dailey et al. 2015; Ji et al. 2019; Layer et al. 2006) are only present in Cj0992c (_14_GPRYTSYPTA_23_ and _311_HRNFQGYTT_319_).

**Figure 3.**
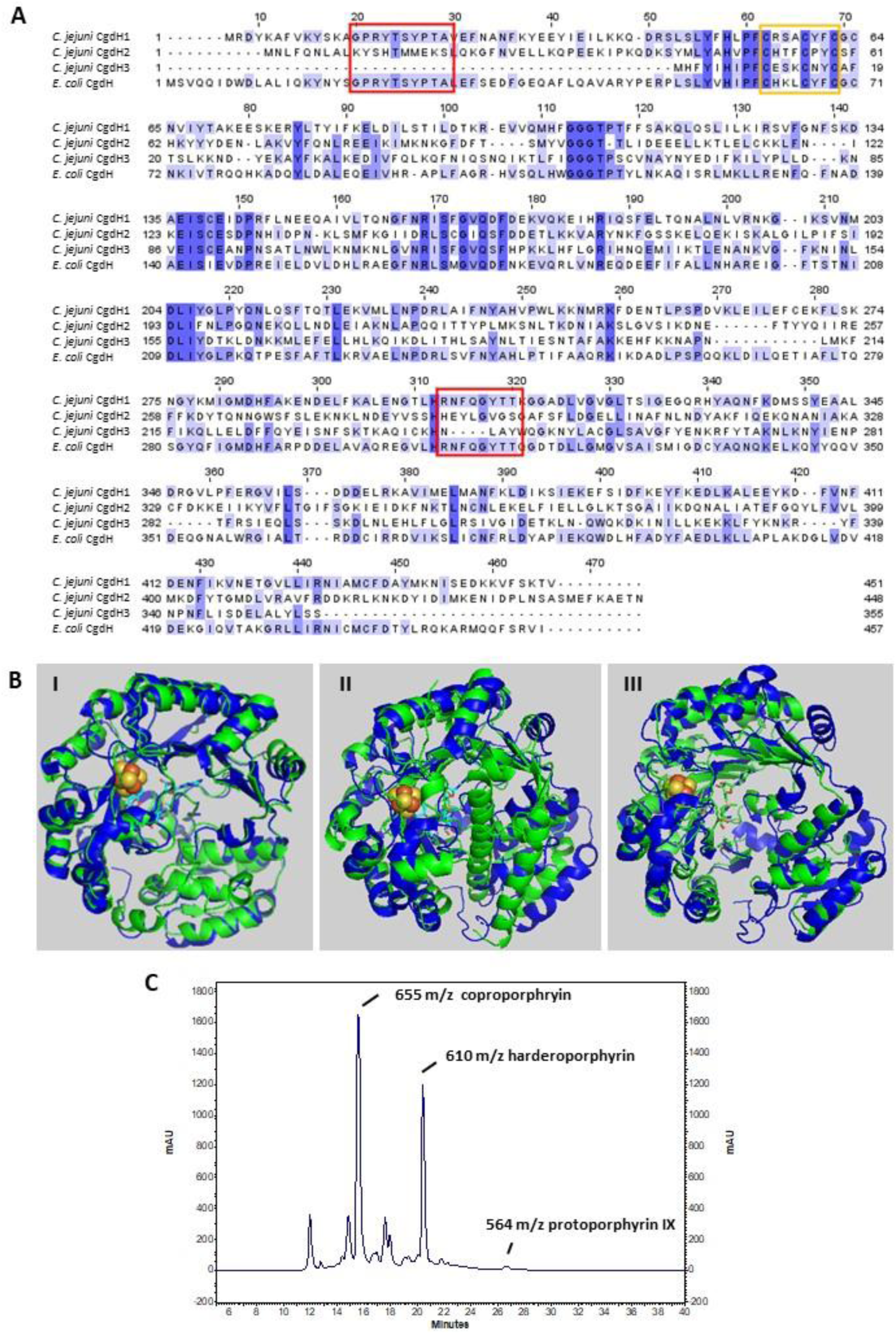
*C. jejuni* CgdH1 has copro’gen dehydrogenase activity. **A.** Amino acid sequence alignment of *C. jejuni* CgdHs, namely CgdH1 (Cj0992c, UniProt (UP): Q0P9R0), CgdH2 (Cj0363c, UP: Q0PBE7), CgdH3 (Cj0580c, UP: Q0PAT7), and CgdH from *E. coli* (UP: P32131), obtained with ClustalOmega and edited with Jalview 2.11.2.5. Dark blue boxes highlight residues with a percentage identity > 80 %. The lighter the blue boxes the lower the percentage identity of the highlighted residues (from 60-0%). Red and orange boxes indicate the motifs required for coproporphyrinogen III dehydrogenase activity (GPRYTSYPTA and RNFQGYTT) and the CxxxCxxC sequence predicted to bind the Fe-S cluster, respectively. **B.** Structures of *C. jejuni* proteins (green), namely CgdH1 (Cj0992c) (I), CgdH2 (Cj0363c) (II) and CgdH3 (Cj0580c) (III), modelled by AlphaFold2 and aligned with *E. coli* CgdH (blue, PDB number 1OLT), with RMSD values of 0.8, 3.3 and 1.7, respectively. **C.** MS spectrum of the protoporphyrinogen IX reaction product of *C. jejuni* Cj0992c. Peaks at m/z 564, 610, 655 correspond to protoporphyrin IX (oxidized form of protógen IX), the intermediate harderoporphyrin I, and the oxidized form of the coproporphyrin III substrate, respectively.

Both Cj0992c and Cj0363c contain putative binding sites for S-adenosyl-L-methionine (SAM) and an Fe-S cluster. Cj0992c contains a typical [4Fe-4S] binding motif (CX_3_CX_2_CXC). In Cj0363c, only 3 of the 4 cysteines are conserved (CX_3_CX_2_C), suggesting that could bind a [3Fe-4S] cluster or that the 4^th^ ligand is not a cysteine and still needs to be identified (Figure 3A).

The *C. jejuni* Cj0992c, Cj0363c and Cj0580c structures were predicted with AlphaFold2 and superimposed with that of *E. coli* CgdH (HemN) (Layer et al. 2003), exhibiting RMSD of 0.8, 3.3 and 1.7, respectively. *C. jejuni* Cj0992c structure is predicted to retain the features of *E. coli* CgdH, i.e., an N-terminal domain significantly larger than the C-terminal domain, which consists of three almost parallel α-helices (Figure 3B-I). On the contrary, the predicted structure of *C. jejuni* Cj0363c is quite different from that of *E. coli* CgdH (HemN; (Layer et al. 2003)), particularly due to the presence of an antiparallel α-helix located between the N-terminal and C-terminal domains (Figure 3B-II). Although the *C. jejuni* Cj0580c modelled structure seems more similar to the *E. coli* CgdH structure than to *C. jejuni* Cj0363c, the alignment of *C. jejuni* Cj0580c with Cj0363c has a RMSD of 1.7 that reflects significant structural differences between the two proteins (Figure 3B-III).

The three *C. jejuni cgdH* genes are spread in the genome. The *cj0992c* gene is part of a gene cluster that also encodes an oxidoreductase ferredoxin-type electron transport protein (Cj0991c), the hypothetical protein Cj0993c, an ornithine carbamoyltransferase ArgF (Cj0994c), and the delta-aminolevulinic acid dehydratase HemB/PbgS (Cj0995c). Genes *cj0363c* and *cj0580c* are scattered throughout the genome and not located near any recognizable heme-biosynthesis related genes.

For sake of simplicity, the three CgdH-like proteins encoded by *cj0992c*, *cj0363c* and *cj0580c* will be designated from now on as CgdH1, CgdH2, and CgdH3, respectively. We cloned, expressed, and purified successfully CgdH1 and CgdH2. However, despite several attempts, the low solubility of CgdH3 (Cj0580c) did not allow obtaining enough quantities for subsequent studies. Although CgdH1-2 contain motifs for binding of Fe-S centers, the proteins were isolated in the apo-form. Hence, prior to the enzyme assays, *C. jejuni* CgdH1 and CgdH2 were treated to promote incorporation of the Fe-S clusters through reconstitution reactions done under anaerobic conditions (see Materials and Methods). After the reconstitution assays, CgdH1 and CghH2 exhibited broad absorption bands typical of the presence of Fe-S centers (∼320 nm and 420 nm) (Supplementary Figure S2).

The copro’gen III dehydrogenase activity of the reconstituted CgdH1 and CgdH2 was evaluated in separate reactions. Each protein was incubated with copro’gen III, SAM and NADH, in the conditions described in Materials and Methods, and the products were analyzed by HPLC-MS. The expected final reactional product protoporphyrin IX (oxidized form of protógen IX) was observed, although the reaction is incomplete as peaks corresponding to the substrate and the harderoporphyrin intermediate are present (Figure 3C). Similar activity assays done for *E. coli* HemN, and analyzed by MS, generated several porphyrin products with the harderoporphyrin intermediate being the highest peak (Rand et al. 2010).

Although the same type of assay was performed with the reconstituted CgdH2, no formation of protoporphyrin IX or any other intermediate was observed, i.e., CgdH2 does not exhibit copro’gen III dehydrogenase activity.

### *C. jejuni* heme pathway: from proto’gen IX to protoporphyrin IX

Three different bacterial enzymes may perform the conversion of proto’gen IX to protoporphyrin IX, namely PgoX, PgdH1 and PgdH2. *C. jejuni* genome search retrieved a PgdH2-like enzyme (Cj0362) that exhibits 39% and 38% identity with PgdH2 from *Synechocystis spp.* and *Acinetobacter baylyi*, respectively (Figure 4A). Despite our efforts to purify *C. jejuni* Cj0362 to homogeneity, it was not possible to obtain enough quantities of the enzyme for the biochemical studies. Alternatively, the proto’gen IX oxidase activity was tested in the membrane fraction of *E. coli* cells transformed with Cj0362 and incubated with the proto’gen IX substrate. Under these conditions, cells expressing Cj0362 exhibited much higher activity for the oxidation of proto’gen IX than cells containing the empty plasmid pPR-IBA2 (Figure 4B). This result indicates that Cj0362 is an active PgdH2 of *C. jejuni*. Since no structures are available for bacterial PgdH2-like proteins, we used AlphaFold2 to model Cj0362, which showed that predicted structure is of the 4-helix-bundled type (Figure 4C).

**Figure 4.**
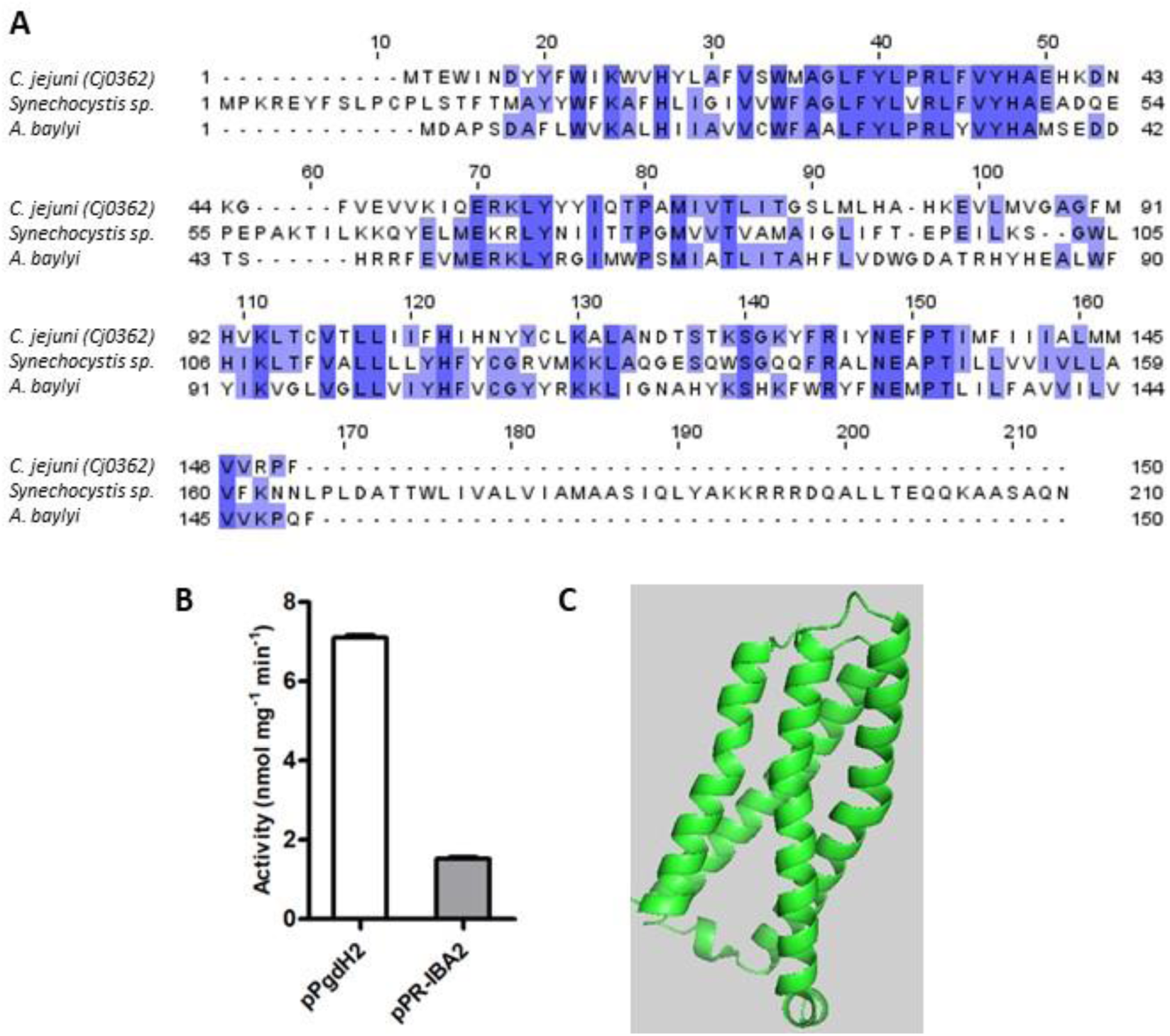
*C. jejuni* contains a proto’gen IX dehydrogenase enzyme. **A.** Amino acid sequence alignment of *C. jejuni* PgdH2 (Cj0362, UniProt (UP): Q0PBE8), with two previously studied protoporphyrinogen oxidases PgoX from *Synechocystis sp.* (UP: P72793) and *Acinetobacter baylyi,* (UP: Q6FDT1), done with ClustalOmega and edited with Jalview 2.11.2.5. Dark blue boxes highlight residues with a percentage identity > 80 %. The lighter the blue boxes the lower the percentage of identity of the highlighted residues (from 60-0%). **B.** Proto’gen IX conversion to protoporphyrin IX was assayed in *E. coli* cell membranes carrying the plasmid expressing *pgdH2* (pPgdH2) or the empty plasmid (pPR-IBA2). The reaction fluorescence of the mixture containing 500 nM proto’gen IX, 150 mM NaCl, 20% glycerol, 5 mM DTT and 0.3 mg of membrane fraction of each *E. coli* cell extract, was monitored for 20 min. The fluorescence intensity units were converted to concentration by interpolation of a calibration curve obtained with protoporphyrin IX standards. **C.** Structure of *C. jejuni* PgdH2 (Cj0362) predicted by AlphaFold2.

### *C. jejuni* heme pathway: from protoporphyrin IX to heme

*C. jejuni* encodes Cj0503c that shares amino acid sequence identity between 22-37 % with ferrochelatase of *Aquifex aeolicus, Rickettsia prowazekii, Human, Mycobacterium tuberculosis,* and *Streptomyces coelicolor* (Figure 5A). Therefore, the recombinant Cj0503c was produced, and the activity of the purified protein was evaluated. *C. jejuni* Cj0503c exhibits a ferrochelatase specific activity of 12.0 ± 1.3 nmol·mg^-1^·min ^-1^, a value that is in the same order of magnitude of other bacterial ferrochelatases (Hansson and Hederstedt 1994; Lobo et al. 2015). Cj0503c activity was also assessed in the heme auxotrophic strain *E. coli ΔppfC* mutant (Miyamoto et al. 1992; Nakahigashi et al. 1991). Complementation of *E. coli ΔppfC* with *cj0503c* allowed growth in heme-free medium, showing that Cj0503c is a functional protoporphyrin ferrochelatase (Figure 5B, Supplementary Figure S1).

**Figure 5.**
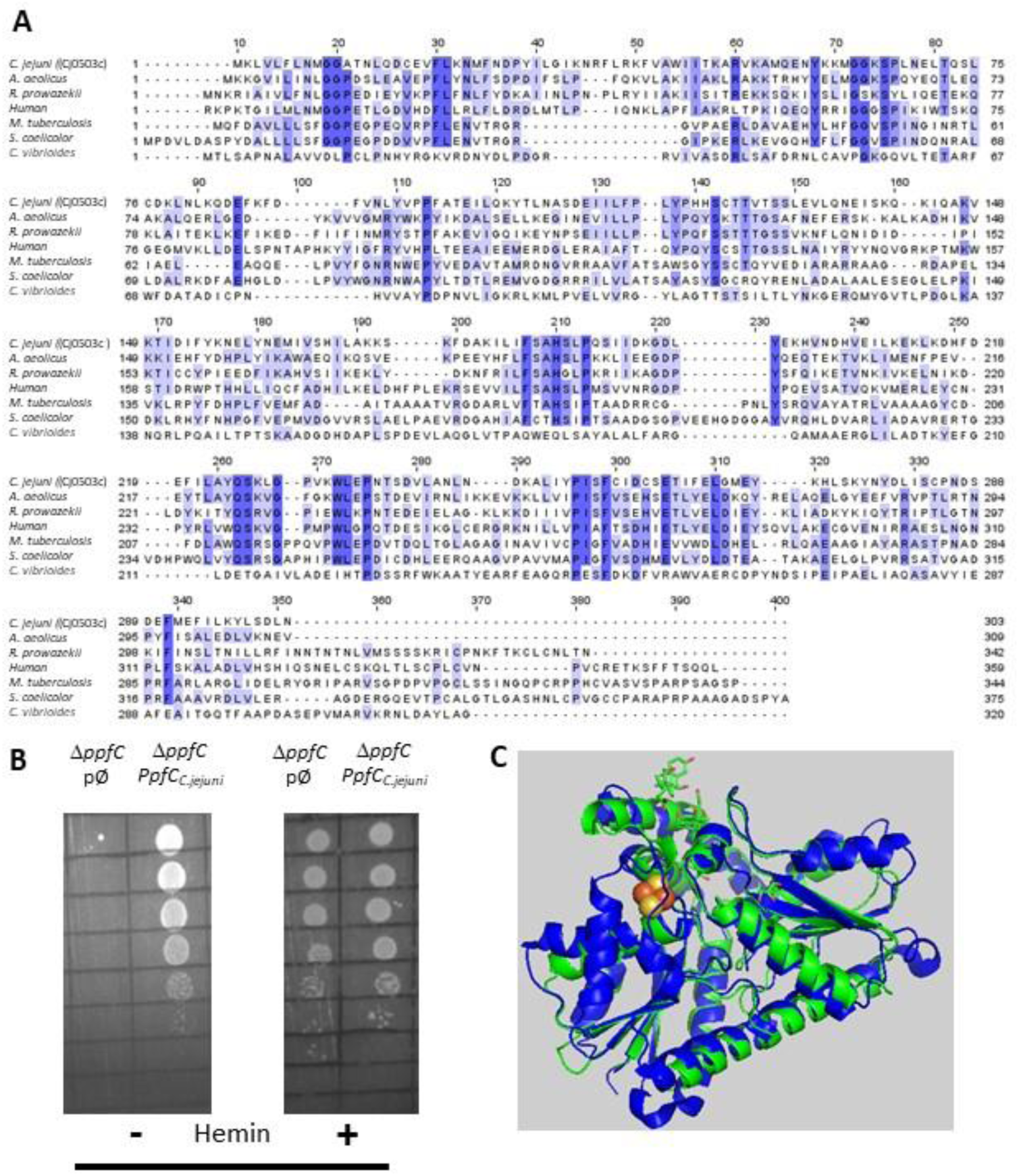
*Campylobacter jejuni* encodes a functional protoporphyrin ferrochelatase (PpfC). **A.** Amino acid sequence alignment of *C. jejuni* PpfC (Cj0503c, UniProt (UP): Q9PI08), and PpfC from *Aquifex aeolicus* (UP: O67083), *Rickettsia prowazek*ii (UP: Q9ZC84), Human (UP: P22830), *Mycobacterium tuberculosis* (UP: P9WNE3), *Streptomyces coelicolor* (UP: O50533) and *Caulobacter vibrioides* (UP: Q9A3G2), done with ClustalOmega and edited with Jalview 2.11.2.5. Dark blue boxes highlight residues with a percentage identity > 80 %. The lighter the blue boxes the lower the percentage identity of the highlighted residues (from 60-0%). **B.** *E. coli* Δ*ppfC* strain transformed with the empty plasmid (pØ) or with a plasmid harboring *C. jejuni cj0503c* gene (*ppfC_C.jejuni_*) was platted in LB-agar medium, in the absence (-) and in the presence (+) of hemin. **C.** Structure modelling using Alphafold2 of *C. jejuni* PpfC (green) superimposed with that of human PpfC enzyme (blue, PDB number 1HRK), with RMSD of 1.2.

The structure of Cj0503c, predicted by Alphfold2, is depicted in Figure 5C. Superimposition of Cj0503c with *B. subtilis* PpfC has a high RMSD of 2.9, but the two structures share a similar overall structure and number of helices and beta sheets.

### *C. jejuni* CgdH2 is a heme binding protein. Identification of the heme-binding residues

We observed that *C. jejuni* CgdH2 has no coproporphyrinogen dehydrogenase activity. A more extensive BLAST analysis showed that CgdH2 shares some degree of similarity with the heme chaperon HemW (e.g, *E. coli* HemW: 24% identity, 49% similarity, 71% coverage) (Figure 6A). This similarity raised the hypothesis that CgdH2 could be a heme binding protein, which was tested in two types of assays.

**Figure 6.**
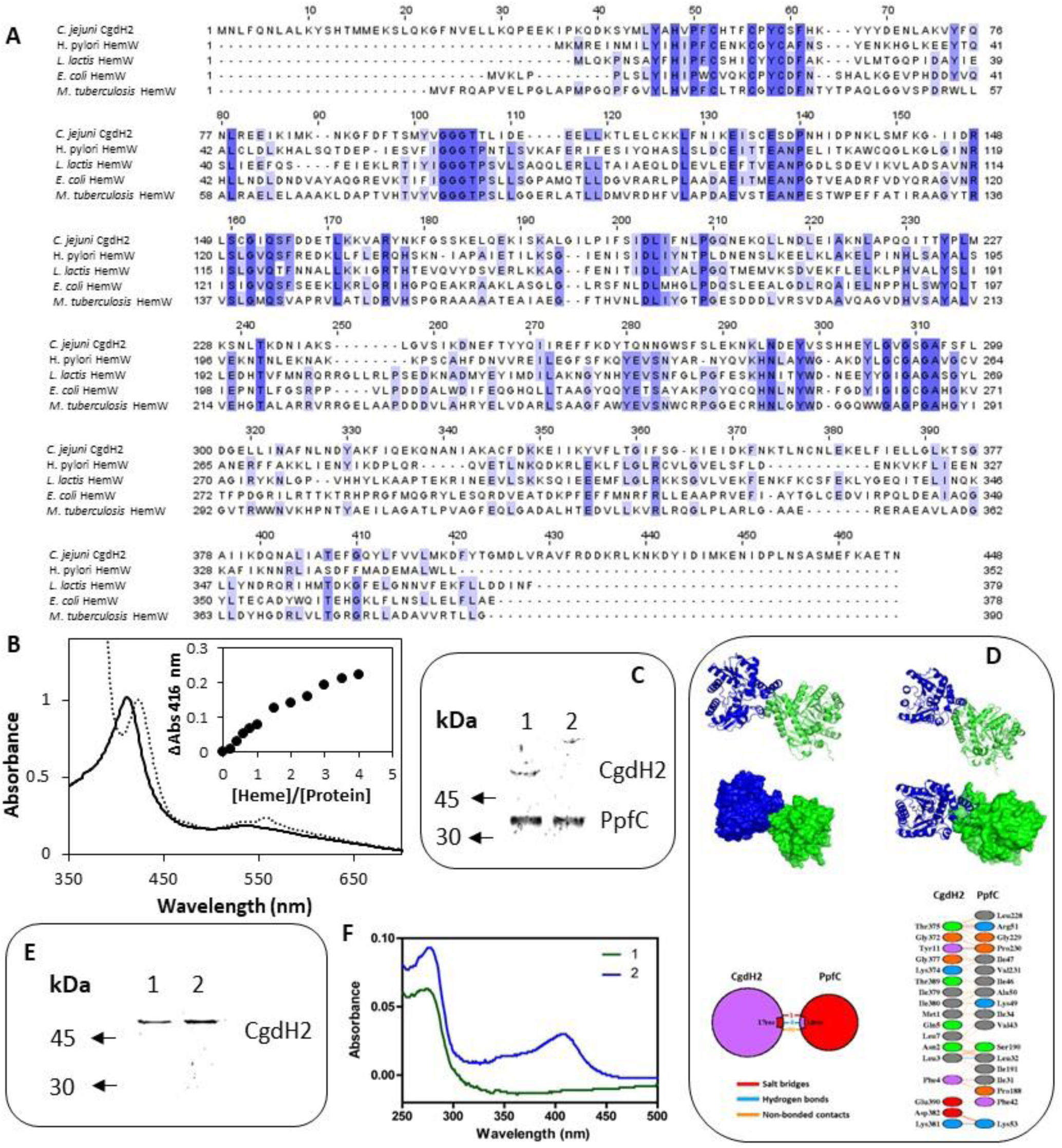
CgdH2 is a heme binding protein that receives heme from ferrochelatase PpfC. **A.** Amino acid sequence alignment of *C. jejuni* CgdH2 (Cj0363c, UniProt (UP): Q0PBE7), and HemW from *Helicobacter pylori* (UP: O25824), *Lactococcus lactis (*UP: Q9CGF7), *E. coli* (UP: P52062) and *Mycobacterium tuberculosis (*UP: P9WP73), done with ClustalOmega and edited with Jalview 2.11.2.5. Dark blue boxes highlight residues with a percentage identity > 80%. The lighter the blue boxes the lower the percentage identity of the highlighted residues (from 60-0%). **B.** Spectroscopic analysis of CgdH2 bound with hemin before ( ) and after reduction with dithionite (----). Inset depicts the titration assay of CgdH2 with increasing concentrations of hemin. ΔAbs represents the absorbance at 416 nm subtracted from the absorbance of the buffer containing only hemin in the corresponding concentration. **C.** Pull-down assay. Apo-PpfC-Strep, apo-CgdH2-His, and a mixture of the two apo-proteins were loaded, separately, onto a Streptactin-Sepharose column. As expected, apo-CgdH2-His does not bind to the column and is eluted with the washing buffer A (Tris-HCl 50 mM pH 8). Fractions eluted with 5 mM desthiobiotin in buffer A were analyzed by SDS-PAGE: lane 1 - mixture of apo-PpfC-Strep with apo-CgdH2-His; and lane 2-apo-PpfC-Strep. **D.** Modelling of the protein interaction between *C. jejuni* PpfC and CgdH2. The docking complex generated by ClusPro 2.0 (Desta et al. 2020; Kozakov et al. 2013, 2017; Vajda et al. 2017) is shown in both ribbon and surface models. The model represents the complex with the lowest energy and higher clustering among the top ten models of the four sets provided by ClusPro. *C. jejuni* PpfC and CgdH2 are represented in green and blue, respectively. Interface residues predicted to be involved in the interaction that were obtained through PDBsum (reference) using the modelled structures are presented in the figure. **E** and **F.** Heme transfer assay. SDS-PAGE (E) and UV-visible spectra (F) of the fractions eluted with buffer A after loading apo-CgdH2 onto an empty resin (1) and onto a resin previously bound to the heme-ferrochelatase (He-PpfC-Strep) (2).

In the first assay, *E. coli* cell extracts expressing CgdH2 were incubated with hemin prior, as described in Materials and Methods. After the purification process, the purified CgdH2 contained one heme molecule (0.8 ± 0.1 heme/protein) as determined by the pyridine hemochromagen assay.

In the second method, *E. coli* cell extracts expressing CgdH2 were not treated with heme to produce apo-CgdH2. The ability of CgdH2 to bind heme was then tested by titrating purified apo-CgdH2 with hemin. Addition of increasing concentrations of hemin to apo-CgdH2 led to an increase in the Soret band at 416 nm (Figure 6B). Data analysis showed that CgdH2 binds one heme molecule per protein, a result that agrees with that obtained in the assay described above. The K_d_ for heme binding of CgdH2 is 4.9 ± 1.0 µM, a value that is in the range of the values described for others heme chaperones, e.g., K_d_∼8 μM for *Lactococcus lactis* HemW (Abicht et al. 2012), K_d_ for Cj1386 of ∼4.3 μM (Flint and Stintzi 2015), and K_d_ of 0.2 μM for *Pseudomonas aeruginosa* heme trafficking protein PhuS (Bhakta and Wilks 2006).

We also tested whether CgdH2 harboring the Fe-S cluster would bind heme. To this end, the Fe-S-reconstitued-CgdH2 protein was incubated with hemin 100 μM and passed through a gel filtration column. The spectrum shows that CgdH2 contains both cofactors, which indicates that the presence of the Fe-S center does not prevent heme binding (Supplementary Figure S2B).

Since we observed that the fully loaded CgdH2 has features resembling a low-spin heme (Figure 6B), we next sought to identify the heme binding ligands by site-directed mutagenesis. We selected ten residues that usually serve as heme ligands, such as histidine, tyrosine, and methionine. The following residues were modified: histidine H48, H53, H62, H133, H285; tyrosine Y46, Y66, Y224, and methionine M44 and M227. The proteins were expressed and cell extracts of each of the single mutants of Cgdh2 were prepared and incubated, separately, with hemin as described above. Following the purification step, the heme content of the purified proteins was determined by the pyridine hemochromogen method. Based on data shown in Table 1, amino acid residues H48 and H133 are predicted to be the heme ligands in CgdH2.

**Table 1.** CgdH2 binds heme. Heme content of CgdH2 wild type and mutant proteins following incubation with hemin, purification, and quantified by the pyridine hemochrome assay. Results are expressed as the ratio of heme per protein concentration.

| CgdH2 protein | Heme/protein | HemW* |
| --- | --- | --- |
| Wild type | 0.8±0.1 | -- |
| **H48L** | 0.3±0.0 | YES |
| H53L | 0.7±0.0 | NO |
| H62L | 0.7±0.0 | NO |
| **H133L** | 0.4±0.0 | NO |
| H285L | 0.8±0.2 | YES |
| Y46L | 0.7±0.1 | YES |
| Y66L | 0.7±0.1 | NO |
| Y224L | 0.8±0.1 | YES |
| M44L | 0.8±0.1 | NO |
| M227L | 0.8±0.2 | NO |

\*Presence of equivalent residues in bacterial HemW proteins, as depicted in Figure 6A; heme ligands of CgdH2 are highlighted in bold.

### 1. *C. jejuni* CgdH2 receives heme from ferrochelatase

In bacteria, there is more than one intracellular pathway for heme *b* formation, but in no case is it known how heme is transferred to target proteins. The ability of *C. jejuni* CgdH2 to bind heme led us to investigate whether the protein could be an intracellular heme receptor. Furthermore, we considered a possible link with enzymes related to heme biogenesis, in particular with ferrochelatase PpfC, which in *C. jejuni* is the last enzyme in the PPD pathway. For this purpose, we constructed, expressed and produced two *C. jejuni* proteins: CgdH2 linked at the C-terminus to a His-tag (apo-CgdH2-His) and PpfC fused at the N-terminus to a Strep-tag (apo-PpfC-Strep). The interaction of the two proteins was analyzed by a pull-down assay. Apo-PpfC-Strep, apo-CgdH2-His, and the mixture of the two apo-proteins, PpfC-Strep with CgdH2-His, were loaded separately onto a Streptactin-Sepharose column that only binds Strep-tagged proteins. Figure 6C shows that apo-CgdH2-His does not bind to the resin and apo-PpfC-Strep is eluted by addition of 5 mM desthiobiotin. However, when the two proteins CgdH2 and PpfC are previously incubated and then loaded onto the resin, CgdH2 is no longer eluted with the washing buffer, but is retained and comes out together with PpfC, as the result of the interaction between the two proteins.

The regions of interaction between the modelled structures of *C. jejuni* PpfC and CgdH2 were predicted using ClusPro and PDBsum. The protein complex includes 16 hydrogen bonds and 4 salt bridges (Figure 6D). Further analysis of the PpfC-CgdH2 complex using PyMol previses additional hydrogen bonds between residues K49 of PpfC and K245 of CgdH2. The main residues of PpfC expected to contribute to the interaction are the positively charged residues R51, K35, K49 and L141. In PpfC, R51 and K49 may bind to N402 and E248 in CgdH2 through a hydrogen bond and a salt bridge contact, respectively (Figure 6D). PpfC K35 may establish several hydrogen bond contacts with different residues of CgdH2, and PpfC K141 can form a hydrogen bond and a salt bridge contact with two different CgdH2 residues (Figure 6D). Moreover, the interaction between PpfC and CgdH2 is predicted to occur in a region where the PpfC heme binding pocket is exposed. This suggests that the heme in PpfC could be transferred throughout this channel to an open region in the structure of CgdH2 (Figure 6D). Therefore, to test whether CgdH2 receives heme from hemin containing PpfC (He-PpfC-Strep), we loaded apo-CgdH2-His onto a Strep-Tactin epharose resin previously bound to He-PpfC-Strep. When the resin was washed with buffer Tris-HCl 50 mM pH 8, the eluted CgdH2 contained heme as judged by SDS-PAGE (Figure 6E) and UV-visible spectroscopy (Figure 6F) as result of the heme transfer from He-PpfC to apo-CgdH2.

### CgdH2 is a novel heme binding protein

We performed cluster analysis for the three CgdH proteins using the Structure Function Linkage Database (SFLD) (Akiva et al. 2014) of the Radical SAM (RSM) proteins superfamily, a subgroup of the coproporphyrinogen III oxidase-like proteins. The results showed that CgdH-like proteins can be divided in 9 functional types, that include the canonical HemN enzymes, HemZ-like, HemW, and ChuW/HutW (Figure 7A). *C. jejuni* CgdH1 is grouped with bona fide HemN enzymes, in agreement with our experimental data. The node containing *C. jejuni* CgdH2 (large blue dot in Figure 7A) clusters with *Gordonibacter* MenK (Uniprot: D6E859; yellow dot in Figure 7A) and other unknow CgdH-like proteins. Importantly, this clade is not neighboring the HemW containing cluster clearly showing that CgdH2 belongs to a different protein family (Figure 7A). We also did a phylogenetic study of the *C. jejuni* CgdH proteins considering the class of CgdH-like proteins (former designated as HemN) that include the bona fide anaerobic coproporphyrinogen III oxidases, methyltransferases, cyclopropanases, and heme chaperones, according to the classification done in Cheng et al. 2022. In brief, 177 sequences belonging to 168 species were used for the phylogenetic reconstruction where 11 clades, corresponding to characterized enzymes, were retrieved (Figure 7B). The *C. jejuni* CgdH1 clusters in the anaerobic coproporphyrinogen III oxidases branch, whereas *C. jejuni* CgdH3 branch in a clade containing CgdH-like proteins of unknown function. Of note, *C. jejuni* CgdH2 is neither in these clades nor in the branch of heme chaperon HemW-like proteins. Instead, CgdH2 appears in the menaquinone methyltransferases branch (Figure 7B), that contains several archaeal MenK proteins. The *C. jejuni* CgdH2 protein is however within a subclade of bacterial MenK sequences where other proteobacterial organisms are also represented.

**Figure 7.**
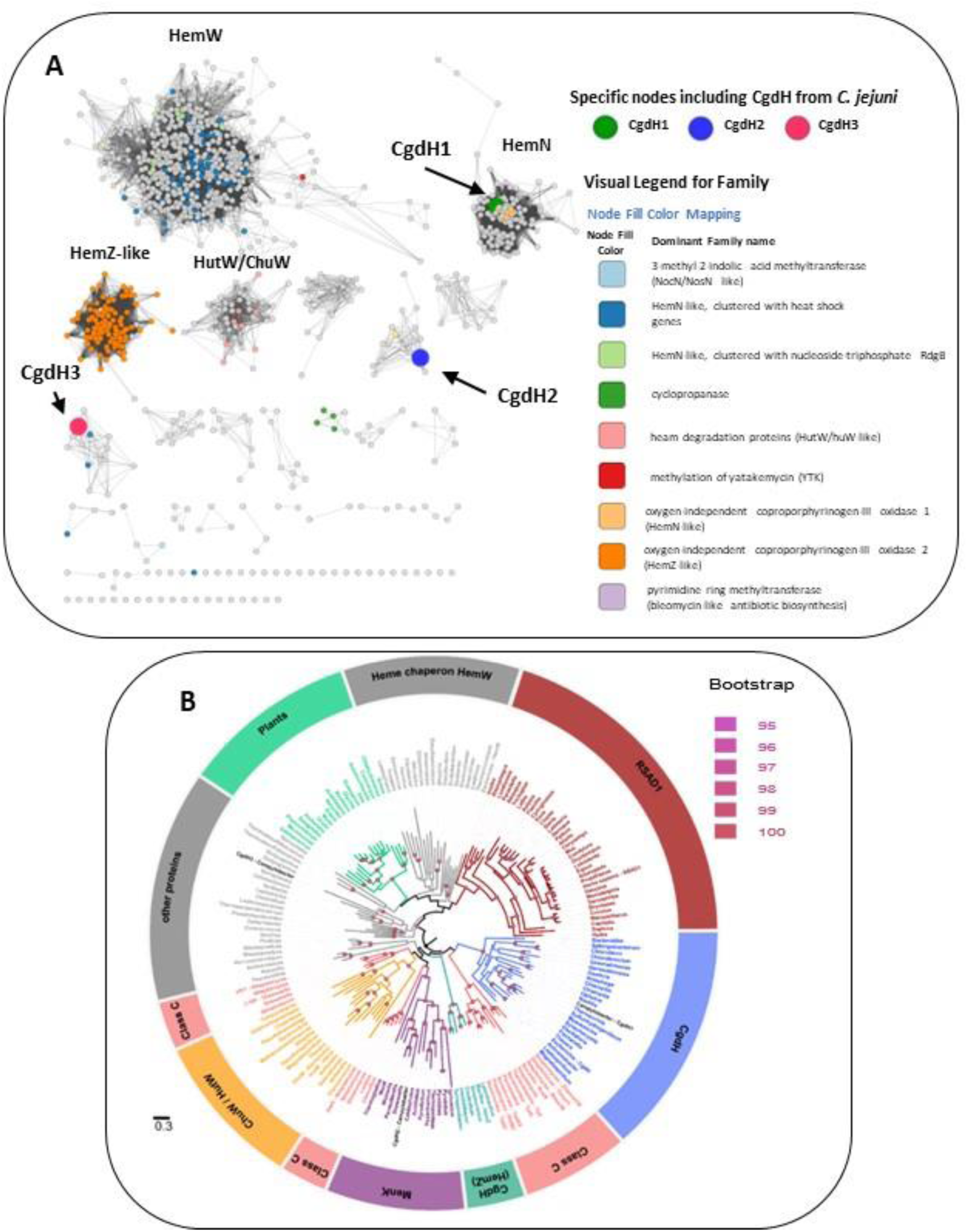
A. Network distribution of the radical S-adenosyl-L-methionine (SAM) anaerobic coproporphyrinogen III oxidase-like enzymes. A. Network distribution obtained using Structure Function Linkage Database (SFLD). Nodes containing CgdH1, CgdH2 and CgdH3 are represented by large green, blue, and red dots, respectively. CgdH1 is included in the bacterial HemNs cluster, while CgdH2 and CgdH3 are located in clusters containing proteins of unknown function. Of note, *Gordonibacter* MenK (CBL03906.1; Uniprot: D6E859) clusters with *C. jejuni* CgdH2 (blue dot) and is represented as yellow dot. B. Maximum likelihood phylogenetic reconstruction of CgdH-like proteins. The phylogeny was rooted using the minimal ancestor deviation method. Ultrafast bootstrap values above 95 (significant) are indicated by filled circles. The different clades are colored based on the functionally characterized CgdH-like protein that is present. The clades obtained were named as: Heme chaperone HemWs, RSAD1 (Radical S-adenosylmethionine domain-containing 1), CgdH (bona fide CgdHs), Class C (class C of Radical SAM methylases), CgdH (HemZ), MenK (menaquinones methyltransferases), ChuW/HutW (heme-degrading enzymes), other proteins (other HemN like proteins), plants (unidentified CgdH-like proteins from plants) (Supplementary Table S4). Genus of the organism containing the CgdH-like protein sequence is indicated. The scale bar indicates the number of substitutions per site.

**Figure 8.**
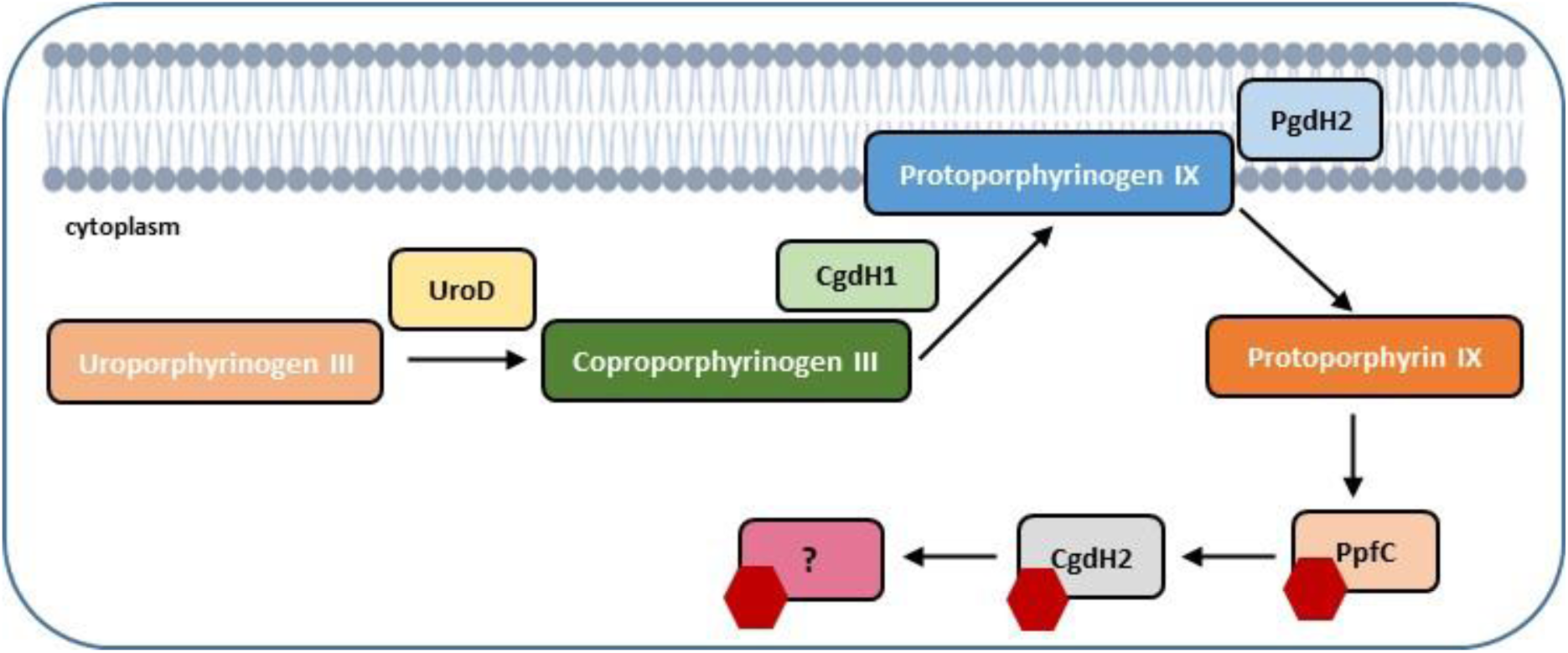
*Campylobacter jejuni* uses the protoporphyrin-dependent (PPD) pathway to form heme. Uroporphyrinogen III decarboxylase (UroD, Cj1243) catalyzes the decarboxylation of uroporphyrinogen III yielding coproporphyrinogen III; oxygen-independent coproporphyrinogen III dehydrogenase (CgdH1, Cj0992c) catalyzes the oxidative decarboxylation of propionate substituents on pyrrole rings A and B of coproporphyrinogen III to the corresponding vinyl groups of protoporphyrinogen IX. In the penultimate step of the pathway, protoporphyrinogen IX is oxidized to protoporphyrin IX by protoporphyrinogen IX dehydrogenase (PgdH2, Cj0362), followed by insertion of ferrous iron into protoporphyrin IX, originating heme (red hexagon) a reaction that is catalyzed by protoporphyrin ferrochelatase (PpfC, Cj0503c). CgdH2 (Cj0363c) receives heme from ferrochelatase (PpfC, Cj0503c) and transfers heme to not yet known target proteins.

Menaquinone methyltransferases include MenK/MenK2/MqnK and belong to the HemN-like class C radical SAM methyltransferases (RSMTs), that share a similar structure build up by a short N-terminal that contains the catalytic domain, a linker domain, and a C-terminal HemN domain. Analysis of a very large number of protein sequences of twelve seed class C RSMTs and 10 other characterized radical SAM proteins (Wilkens et al. 2021). The eight main clusters were designated as HemW, HemZ, ChuW, HutW, MqnK/MenK/MenK2 (MKMT), Jaw5, NosN, and C10P. Six signature motifs were identified and their occurrence allowed defining the HemW and MKMT subgroups (Wilkens et al. 2021). Here, we analyzed the presence of these motifs in CgdH2 when compared with those present in HemW and MKMT proteins (Table 2). Table 2 shows that the PFCxxCxxCF motif, that ligates the [3Fe-4S] center, is conserved in all proteins. The two motifs YFxxxRxE and QxTxYPLx that are absent in HemWs occur in CgdH2. Motifs Y/VxGGGT and NxFxxxxY that have very low occurrence in HemW (∼2%) are present in CgdH2. Moreover, CgdH2 does not contain the H_250_NxxYW_255_ motif (*E. coli* HemW numbering) that is conserved in all bacterial HemW (Dailey et al. 2015) (Figure 3A). In addition, superimposition of the predicted structures of *E. coli* HemW and CgdH2 shows a non-significant RMSD of 2.6. Altogether, we concluded that CgdH2 is not a HemW-like protein but constitutes a novel type of heme chaperone.

**Table 2.**
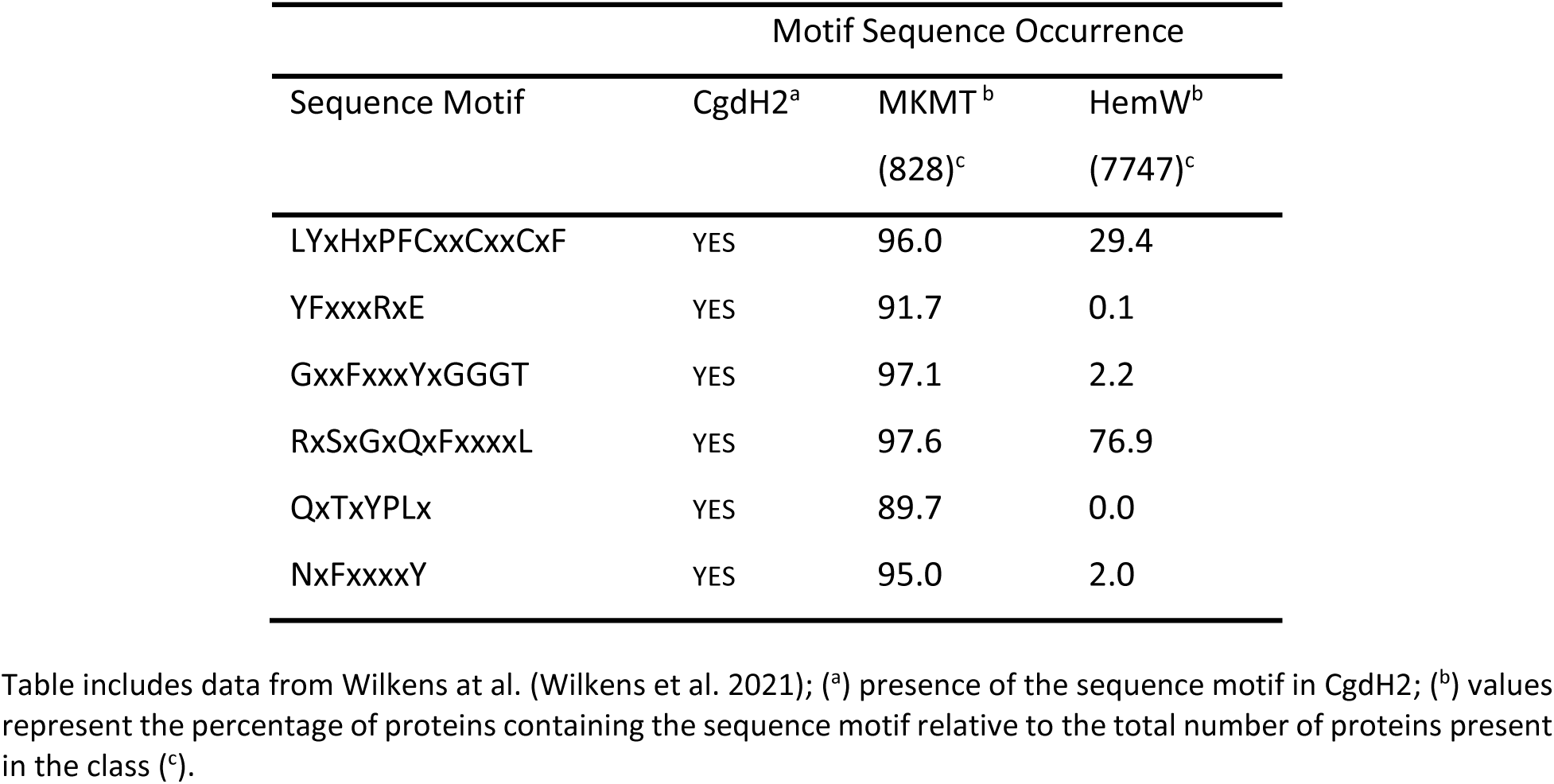
Occurrence of specific sequence motifs in CgdH2, menaquinone methyltransferases (MKMT), and HemW-like chaperones.

## Discussion

Bacterial heme biosynthesis begins with the formation of 5-aminolevulinic acid that is converted to the intermediate tetrapyrrole macrocyclic uro’gen III. The biochemical characterization of several enzymes allowed us to show that in Gram-negative *C. jejuni* the subsequent transformation of uro’gen III into heme occurs through a PPD-like pathway (Figure 1). Furthermore, we clarified which of the three putative copro’gen dehydrogenase proteins retrieved by BLAST search analysis of the *C. jejuni* genome is the enzyme acting in the heme biogenesis pathway.

The heme formed in ferrochelatase through the biogenesis route needs to be escorted by protein chaperones to the final destination in a highly controlled manner. Several proteins may play this role based on their heme-binding affinity, such as the group of chaperone-like monoheme protein HemWs (Haskamp et al. 2018). The *E. coli* HemW exhibits a weak SAM cleavage activity and harbors an [4Fe–4S] that although not required for heme binding is necessary for heme transfer to the membrane subunit NarI of the respiratory nitrate reductase NarGHI. And the transfer of heme from HemW to a catalytically inactive enzyme NarI restores the nitrate reductase activity*. E. coli* HemW interacts with NarI, but it does not interact with ferrochelatase, bacterioferritins BfrA and BfrB, and catalase KatA (Haskamp et al. 2018).

Another type of heme chaperones are the mycobacterial orthologues which were shown, by phylogenetic analysis, to form a divergent clade that includes the HemW proteins of other species such as *Escherichia coli* and *Lactococcus lactis* (Sharma et al. 2021). Molecular dynamics studies showed that in mycobacteria HemW binds heme through the conserved H_250_NXXYW_255_ motif. In *C. jejuni* CgdH2 this motif is absent, and while H45 is highly conserved in all HemW-like proteins, residue H133 is unique to CgdH2 (Figure 5A). Moreover, ten single point mutation variants revealed that histidine residues H45 and H133 are required to bind and/or to stabilize the heme ligation.

Cj1386 is a heme binding chaperon previously described in *C. jejuni*, which binds one heme coordinated through tyrosine Y57 (Flint and Stintzi 2015). The decreased hemin content of KatA in the Δ*cj1386* mutant was proposed to account for the reduced catalase activity and, consequently, the higher sensitivity of the strain to oxidative stress. Therefore, Cj1386 is considered a chaperon that binds and traffics heme to KatA (Flint and Stintzi 2015). *C. jejuni* CgdH2 (447 aa) and Cj1386 (156 aa) share a low sequence similarity (I <30%, query coverage of 18%). Furthermore, *C. jejuni* Δ*cgdH2* mutant has no modified sensitivity to oxidative stress when compared with the wildtype strain and CgdH2 does not interact with KatA (data not shown).

The interaction between ferrochelatases and other enzymes of the heme biosynthesis pathway has been previously reported as following described. In the cyanobacterium *Thermosynechococcus elongatus* ferrochelatase interacts with proto’gen IX oxidase; in *Vibrio vulnificus* a complex between ferrochelatase and proto’gen IX dehydrogenase was described; in *B. subtilis*, CpfC ferrochelatase co-purifies with Fra, a homolog of the eukaryotic iron chaperon frataxin; and in *S. aureus* there is a transient interaction between ferrochelatase and coproheme decarboxylase ChdC. Also in *S. aureus*, ferrochelatase CpfC was shown to interact with heme oxygenase IsdG of the heme uptake system (Reniere, Haley, and Skaar 2011; Videira et al. 2018; Zamarreño Beas et al. 2022). In this work, we provide evidence for the interaction of ferrochelatase and transfer of heme to a novel type of heme chaperone. *C. jejuni* CgdH2 belongs to the radical SAM superfamily, which includes the canonical anaerobic coproporphyrinogen III oxidase-like enzymes, heme oxygenase and heme chaperones. However, CgdH2 falls out of the HemW containing clusters as shown by the SFLD and phylogenetic analysis (Figure 7). Interestingly, CgdH2 appears in the menaquinone methyltransferases-like proteins (MKMT) clade due to the conservation of several amino acid sequence motifs. The homology between sequence and structural features of CgdH2 and MKMTs (e.g., RMSD of ∼0.89 between *C. jejuni* CgdH2 and *Gordonibacter pamelaeae* MenK) also raises the possibility of the involvement of CgdH2 in methylation reactions, a hypothesis that needs to be addressed in future studies.

## Conclusion

Altogether, our biochemical and phylogeny distribution data show that *C. jejuni* CgdH2 is a novel type of heme chaperone, with features that differ from that of bacterial HemWs. Moreover, we show that CgdH2 accepts the heme formed in the PPD tetrapyrrole biosynthesis pathway.

The existence in *C. jejuni* of at least two heme chaperones also shows that heme homeostasis requires multiple chaperones with specific functions, and the data herein gather enlarges our knowledge of the players that are involved in the intracellular binding/transporter of heme.

### Experimental procedures

#### Growth of strains and gene cloning

The strains used in this work are listed in Supplementary Table S1. *E. coli* strains were grown under aerobic conditions in Luria-Bertani (Roth) broth, at 37 °C and 150 rpm. *C. jejuni* NCTC 11168 strain was first cultured in Muller-Hinton agar plates and then grown in Muller Hinton broth in flasks filled with 1/10 of the volume and incubated in a microaerobic environment (83.3% N_2_, 7.1% CO_2_, 3.6% H_2_, 6% O_2_). Cultures were supplied with the appropriate antibiotics, such as ampicillin (50 µg·mL^-1^), kanamycin (50 µg·mL^-1^), and chloramphenicol (30 µg·mL^-1^)

1. *C. jejuni* NCTC 11168 genes encoding UroD (*cj1243*), CgdH1 (*cj0992c*), CgdH2 (*cj0363c*), CgdH3(*cj0580c*), PgdH2 (*cj0362*), and PpfC (*cj0503c*) were amplified from genomic DNA extracted according to manufacturer’s instructions (QIAprep Spin Miniprep Kit) using the by standard PCR reactions containing Phusion High-Fidelity DNA Polymerase (Thermo Fischer) and the primers described in Supplementary Table S2. DNA fragments were cloned into the desired plasmid by digestion and ligation by T4 DNA ligase.

We used pPR-IBA2 vector (IBA) to produce the N-terminal Strep-tag fused proteins and pET-23b for C-terminal His-tag fused proteins. The correct plasmid insertions were confirmed by DNA sequencing. Plasmids pPR-IBA2 containing *C. jejuni uroD* or *C. jejuni ppfC* were used in complementation assays. *E. coli* Δ*ppfC* and *E. coli* Δ*uroD* mutant strains were obtained from National BioResource Project (NBRP) at National Institute of Genetics (NIG), https://shigen.nig.ac.jp/ecoli/strain/. These strains were made competent and transformed by heat shock treatment (42 °C) with plasmids expressing *C. jejuni* PpfC and *C. jejuni* UroD. Cells were grown overnight in LB supplemented with hemin (10 mM). After ∼16h of growth, 1 mL of the overnight culture was centrifuged at 1,700 x *g* and resuspended 3 times in PBS and diluted 1:10 six times. From each dilution, 5 mL were spotted in LB agar plates supplemented, or not, with hemin in the indicated concentrations. The strains, plasmids and fusion proteins used in this study are presented in Supplementary Table S3.

#### Production and purification of recombinant proteins

Cells of BL21STAR(DE3)pLysS (Novagen) transformed with pPR-IBA2-*uroD-strep* plasmid, pET23b-*cgdH1* and pET23b-*cgdH2-his* were grown separately overnight, diluted 1:100 times in fresh LB containing ampicillin. The ratio 2/5 of liquid medium/flask volume was used. Cells were grown at 37 °C 150 rpm to an OD∼0.6. At this point, isopropyl β-D-1-thiogalactopyranoside (IPTG) induced expression of UroD (50 µM IPTG), CgdH1 and CgdH2 (500 µM IPTG). The temperature was lowered to 20 °C, and cells grown for 16-20 h. Cells were harvested by centrifugation (11,000 x *g*, 10 min, 4 °C), and pellets were resuspended in 100 mM Tris-HCl pH 8 (buffer A) with 150 mM NaCl. Cells were disrupted in a French Press operating at 1,000 Psi and centrifuged (42,000 x *g*, 40 min, 4 °C).

For purification of *C. jejuni* UroD-Strep, the supernatant was applied onto Strep-Tactin Superflow® high-capacity resin (Iba®), previously equilibrated with buffer A, through a 0.45 mm filter. After washing the column with at least 3 column volumes of buffer A, proteins were eluted with buffer A containing 150 mM NaCl and 2.5 mM desthiobiotin. After elution, all further steps were done under anaerobic conditions. A buffer exchange to Tris-HCl 50 mM pH 8 (buffer B) was done in a PD-10 column, and the protein was concentrated in a diaflow with suitable cut-off membranes (EMD Millipore Corporation, Billerica, Massachusetts).

For *C. jejuni* CgdH1-His and CgdH2-His purification, the respective supernatants were applied, separately, into Chelating Sepharose™ fast flow resin columns (GE Healthcare, Carnaxide, Portugal) charged with Ni^2+^ and previously equilibrated with 4 column volumes of buffer B plus 500 mM NaCl and 10 mM imidazole. After washing the columns with buffer B containing 500 mM NaCl, buffers with increasing concentrations of imidazole were applied (10 mM - 500 mM), and His-tag CgdH1 and CgdH2 proteins were eluted at 400 mM imidazole.

For production of *C. jejuni* PpfC-Strep enzyme, 1% of a pre-culture of *E. coli* BL21STAR(DE3)pLysS (Novagen) cells harboring the pPRIBA2-*ppfC*-*strep* was used to inoculate fresh LB media containing ampicillin. When cells reached an OD_600_ ∼ 0.7, 500 μM of IPTG was added and the cells further grown for 3.5 h, at 30°C at 150 rpm. Cells were harvested by centrifugation (11,000 × *g*, 10 min, 4 °C), and the pellets were resuspended in buffer B supplemented with 150 mM NaCl, 10% glycerol and 1% Triton X-100. Cells were disrupted in a French press operating at 1,000 Psi, and centrifugated at 48,000 × *g*, at 4 °C, for 30 min. The lysates were added to a Strep-Tactin Sepharose column washed with buffer B supplemented with 400 mM NaCl, 10% glycerol, and with increasing concentrations of desthiobiotin. The protein was eluted with buffer B - 5 mM desthiobiotin. All protein fractions were dialyzed in buffer B with 100 mM NaCl and its purity confirmed by SDS-PAGE.

For the iron-sulfur (Fe-S) center reconstitution assays of the purified *C. jejuni* His-tagged CgdH1 and CgdH2, the proteins were diluted to 200 µM in buffer B with 5 mM DTT, 150mM NaCl and 20% glycerol followed by addition of eight-fold molar excess of FeCl_3_. After 5 min incubation, lithium sulfide was added at the same molar concentration as iron, and the mixture was incubated at 4 °C for 3 h. Unbound iron and sulfide were removed using a PD-10 desalting column (GE Healthcare). Formation of the Fe-S center was confirmed by UV-visible spectroscopy (Supplementary Figure S3).

#### Biochemical assays

Porphobilinogen, coproporphyrin III, protoporphyrin IX and hemin were purchased from Frontier Scientific. Freshly prepared enzymes were always used, except if otherwise indicated, and assays were in a Coy model A-2463 Belle Technology anaerobic chamber containing a Shimadzu UV-1800 spectrophotometer. Porphyrins were detected by their spectral absorbance bands localized 390 nm and 410 nm. Unless otherwise indicated, reactions were done in buffer B, at room temperature.

*C. jejuni* UroD activity was determined by a linked protein assay that allow the production of uroporphyrinogen III (uro’gen III), as described previously (Lobo et al. 2015). Briefly, reaction mixtures done in buffer B (final volume of 2 mL) containing 2 mg of *Desufovibrio (D.) vulgaris* PbgS, 2 mg of *D. vulgaris* UroS, and 1.5 mg of porphobilinogen were prepared in the absence (control reaction) and in the presence of 1.5 mg of *C. jejuni* UroD, incubated overnight and oxidized by addition of 0.1 M HCl. Following a centrifugation step (10,000 x g, 10 min) to remove protein precipitate, the formation of the substrate uro’gen III (present in the control reaction) and product copro’gen III (resultant from the activity of UroD) were detected in their oxidized form, namely as uroporphyrin III and coproporphyrin III, by UV-visible spectroscopy and HPLC-MS analysis. For the later, samples were resolved by HPLC-MS on an Ace5-AQ column attached to an Agilent 1100 series HPLC equipped with a diode array detector and coupled to a micrOTOF-Q II (Bruker) mass spectrometer. Products separation was achieved by applying a binary gradient of 0.1% TFA (solvent A) and acetonitrile (solvent B), at a flow rate of 0.2 mL·min^−1^. Column was equilibrated with 80% of solvent A and 20% of solvent B, the samples were injected and concentration gradient of solvent B reached 100% in 50 min.

For *C. jejuni* CgdH1 and CgdH2 activity assays, copro’gen III was first prepared in a reaction mixture containing *D. vulgaris* PbgS, *D. vulgaris* UroS, porphobilinogen, and *C. jejuni* UroD in concentrations like those described above. The concentration of copro’gen III formed was determined after addition of 0.1 M HCl spectroscopically at 548 nm (ε_548_ = 16.8 mM^−1^·cm^−1^). In separated reactions, each *C. jejuni* CgdH with Fe-S centre reconstituted (∼20 μM) was incubated with 10 μM of copro’gen III, 1 mM of SAM, 0.5 mM of NADH, and 30 μL *E. coli* BL21STAR(DE3)pLysS cell extract (10 mg/mL of protein concentration). Reaction mixture was incubated overnight at room temperature in the anaerobic glove box. The resultant samples were treated with 0.6 M HCl and centrifuged at 10000 × g, for 10 min, to remove protein precipitates. The porphyrins formed in the reaction were separated in an Ace5-AQ C18 column, attached to a Hitachi LaChrom Elite HPLC equipped with a diode array detector (VWR, Alfragide, Portugal), and using the same solvents and gradients above described.

Proto’gen IX was prepared by reduction of protoporphyrin IX using a sodium mercury amalgam. Approximately 3 mg of protoporphyrin IX was solubilized with three drops of 30% ammonium hydroxide and dissolved in 5 mL of freshly prepared 10 mM KOH. Concentration was measured and the porphyrin was diluted to 50 μM in a 2 mL final volume. After addition of 5% sodium mercury amalgam (Sigma), the flask was sealed, degassed with nitrogen, and 100 μL of 1 M Tris-HCl pH 8 was added, and the reaction kept in the anaerobic chamber protected from light. The solution containing proto’gen IX was filtered and the concentration of the compound was determined spectroscopically in 0.1 M HCl (ɛ_408n_m= 297 mM^-1^ cm^-1^).

For determination of proto’gen IX oxidase activity, we prepared reaction mixtures in buffer B (final volume 200 μL) containing 500 nM proto’gen IX, 150 mM NaCl, 20% glycerol, 5 mM DTT, and 0.3 mg of the membrane fraction of *E. coli* BL21STAR(DE3)pLysS cell extracts either expressing *C. jejuni* PgdH2 or the empty pPR-IBA 2 plasmid (negative control). Samples were prepared in black 96 well plates (Greiner) and the fluorescence was monitored for 20 min. The formation of protoporphyrin IX was evaluated using a 390 nm excitation filter and 630 nm emission filter in a Fluorostar Optima plate reader (BMG Laboratories), under aerobic conditions. The fluorescence intensity units were converted to concentration by interpolation of a calibration curve obtained with protoporphyrin IX standards. The auto-oxidation of the substrate was subtracted from the oxidation generated by the membrane fractions. The initial rate was calculated from the linear region of the graph.

For the ferrochelatase activity, protoporphyrin IX was prepared as described previously (Lobo et al. 2015). Briefly, ∼1 mg of the compound was solubilized in 1-3 drops of 25% NH_4_OH and subsequently diluted in 500 μL Triton X-100 and 4.5 mL water. The concentration of protoporphyrin IX was determined spectrophotometrically in 2.6 M HCl at 408 nm (ε=297 mM^−1^cm^−1^). Reaction mixtures contained 10 μM of protoporphyrin IX and 30 μg of *C. jejuni* PpfC in buffer B supplemented with 20 μM of (NH4)_2_Fe(SO4)_2_, 5 mM DTT, 150 mM NaCl and 20% glycerol. *C. jejuni* PpfC activity was assayed by following the decrease of the amount of the protoporphyrin IX substrate monitored at 408 nm.

#### Heme binding, pull down, and heme transfer assays

For the heme binding assay, a soluble fraction expressing CgdH2 (prepared as described above) was incubated with 300 μM of hemin (Roth), for 1 h at room temperature. The protein solution was loaded onto a Chelating Sepharose™ fast flow column (GE Healthcare) equilibrated with 20 mM Tris-HCl pH 8 containing 500 mM NaCl and 10 mM of imidazole. The column was washed with increasing concentrations of imidazole, and the protein was eluted with 0.5 M of imidazole. To remove any unbound hemin, the pure protein was loaded onto a PD-10 column (GE Healthcare) that was washed with 20 mM Tris-HCl pH 8 buffer. The heme bound to the protein was quantified by measuring the absorption spectra of pyridine hemochromes (Berry and Trumpower 1987).

The heme titration experiment was performed essentially as described previously (Lobo et al. 2017). Briefly, purified *C. jejuni* apo-CgdH2 was diluted to final concentration of 5 μM in buffer B and loaded into a magnetically stirred quartz (10 mm) cuvette. Increasing concentrations of hemin (from 1 to 20 μM), diluted in 0.1 M of NaOH, were added to the protein solution. The procedure was also done using a similar solution mixture that did not contain protein (control reaction). The titration was monitored by UV-visible spectroscopy in a Shimadzu UV-1700 spectrophotometer, at room temperature. The binding stoichiometry and affinities were determined by plotting the absorbance difference spectra measured at 416 nm.

For the pulldown assay, apo-PpfC-Strep was incubated for 30 min with apo-CgdH2-His, in buffer B, at 4°C. The apo-PpfC-Strep, apo-CgdH2-His, and a mixture of apo-PpfC-Strep with apo-CgdH2-His were loaded, separately, onto Strep-Tactin Sepharose resin containing columns. The resins were washed with 3 volumes of buffer B containing 400 mM NaCl and 10% glycerol (buffer B), 2 volumes of buffer Bsalt supplemented with 5 mM desthiobiotin.

For the heme transfer assays, a cell lysate expressing *C. jejuni* PpfC-Strep was prepared as described above. Prior to the purification step, the cell lysate was incubated with ∼100 μM of hemin for 1 h at room temperature, after which was loaded onto the Strep-Tactin Sepharose column and purification followed the above indicated protocol. The purified PpfC was isolated bound to heme and designated as He-PpfC-Strep. To test the heme transfer between the *C. jejuni* proteins, we mixed purified He-PpfC-Strep with purified apo-CgdH2-His for 1 h at room temperature in buffer B. The apo-CgdH2-His, He-PpfC-Strep, and the mixture of the two after incubation were loaded, separately, onto Strep-Tactin® Sepharose resins. The resins were washed with buffer B containing 400 mM NaCl and 10% glycerol with or without supplementation of 5 mM of desthiobiotin. Fractions were analyzed by SDS-PAGE and visible spectroscopy.

#### Bioinformatic and phylogeny analysis

Basic local alignment search tool (BLAST) was used to search genomes for proteins involved in tetrapyrrole biosynthesis, and sequence alignments were done with ClustalW using the predefined parameters.

The three dimensional protein structures were predicted using the software AlphaFold2 through MMseqs2 and its predefined settings (Mirdita et al. 2022). The similarity between the structures was determined through the PyMOL software (Schrödinger 2020). Data quality was evaluated through the Root Mean Square Deviation (RMSD) value that reflects the average distance between the atoms of each protein structure, with values decreasing with the increasing fitting between the two structures. Structures were visualized using PyMOL and used for in silico protein docking studies.

The predicted interacting regions of *C. jejuni* PpfC and CgdH2 were obtained using the modelled structures of the proteins in molecular docking ClusPro 2.0 server (Desta et al. 2020). The program evaluates multiple number of conformations retaining the most favorable interactions based on thermodynamically energy calculations and provides a list of balanced, electrostatic-favored, hydrophobic favored and Van der Walls electrostatic-fevered interactions (Lensink and Wodak 2013). Interacting residues on *C. jejuni* PpfC-CgdH2 protein complex were analyzed with PDBsum generate (Laskowski 2001) and visualized using PyMOL.

The subgroup of the Radical SAM superfamily that includes the anaerobic coproporphyrinogen-III oxidase-like enzymes from the Structure Function Linkage Database network (repnet.sg1065.th50.pE20.mek250.xgmml) (Akiva et al. 2014) was used in Cytoscape (v3.8.0; following the protocol described in www.sfld.rbvi.ucsf.edu to distinguish the three CgdH proteins. This network distribution is based on the enzyme sequence, structure and molecular function. The database includes cyclopropanases, heme degradation proteins (HutW/ChuW-like), HemN-like clustered with heat shock encoded genes, HemN-like clustered with nucleoside-triphosphate RdgB, oxygen-independent coproporphyrinogen-III oxidase 1 and 2 (HemN and HemZ-like). The nodes were organized based on the ‘rep-net mean’ value, and nodes with a - log_10_(E-value) below 60 are not considered in the analysis.

We performed a cluster analysis for the three CgdH proteins using the Structure Function Linkage Database (SFLD) of the Radical SAM (RSM) proteins superfamily, subgroup of the coproporphyrinogen III oxidase-like proteins (1045 nodes, ∼65000 protein GIs) and each node represents a set of proteins that share at least 50% identity, i.e, it is a group of similar sequences (a total of ∼100 nodes). Edges between nodes indicate a mean BLAST E-value, among all pair sequences in these nodes with an e-value lower than 1e-60 (Akiva et al. 2014).

For the phylogeny analysis, protein sequences selected by Cheng et coworkers (Cheng et al. 2022) were retrieved from NCBI database based on accession number and mapped to NCBI Taxonomy. CgdH1, CgdH2 and CgdH3 sequences were also included in the analysis (Supplementary Table S4). A multiple sequence alignment was performed with ClustalOmega (https://www.ebi.ac.uk/Tools/msa/clustalo/ (Sievers et al. 2011) with: Max guide tree operations = 5, Max HMM iterations = 5, and then trimmed with trimAl v1.3, (Capella-Gutiérrez, Silla-Martínez, and Gabaldón 2009)) through Phylemon 2 webserver using as parameters: Minimum percentage of positions to conserve = 60, gap threshold = 0.05. A maximum likelihood phylogenetic reconstruction was done in IQ-TREE webserver (Trifinopoulos et al. 2016) considering 1000 ultrafast bootstrap and best model selection. The phylogenetic reconstruction was rooted using the minimal ancestor deviation (MAD) method (version 2.22, (Tria, Landan, and Dagan 2017)) with a modified script to keep bootstrap values. FigTree was used for annotations and analysis (v1.4.4, http://tree.bio.ed.ac.uk/software/figtree/).

## Supplementary Material

**Figure S1.**
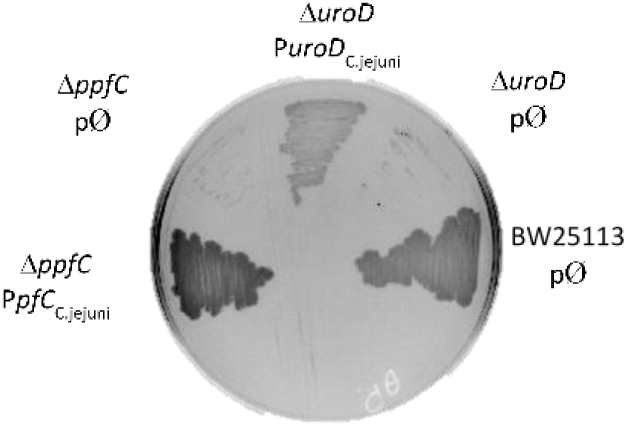
Functional complementation by *C. jejuni* UroD and PpfC enzymes. *Escherichia coli* wild type, Δ*uroD*, aqnd Δ*ppfC* strains expressing the empty plasmid (pØ) or a plasmid containing a functional copy of the correspondent *C. jejuni* homologous gene (P*uroD_C.jejuni_* or P*pfC_C.jejuni_*) streaked in LB-agar medium with no hemin.

**Figure S2.**
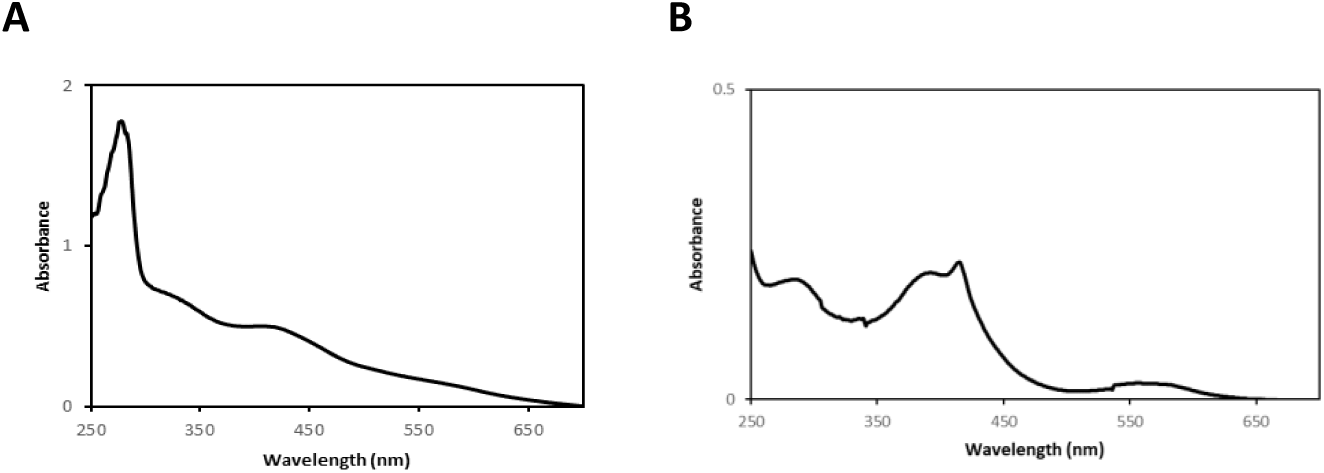
UV-Visible spectra of *C. jejuni* CgdHs proteins. **A.** CgdH1 spectrum exhibits two broad bands centered around 330 nm and 420 nm characteristic of a [4Fe-4S]2+ cluster. **B.** The spectrum of *C. jejuni* CgdH2 indicates the presence of an Fe-S center and heme.

**Table S1.**
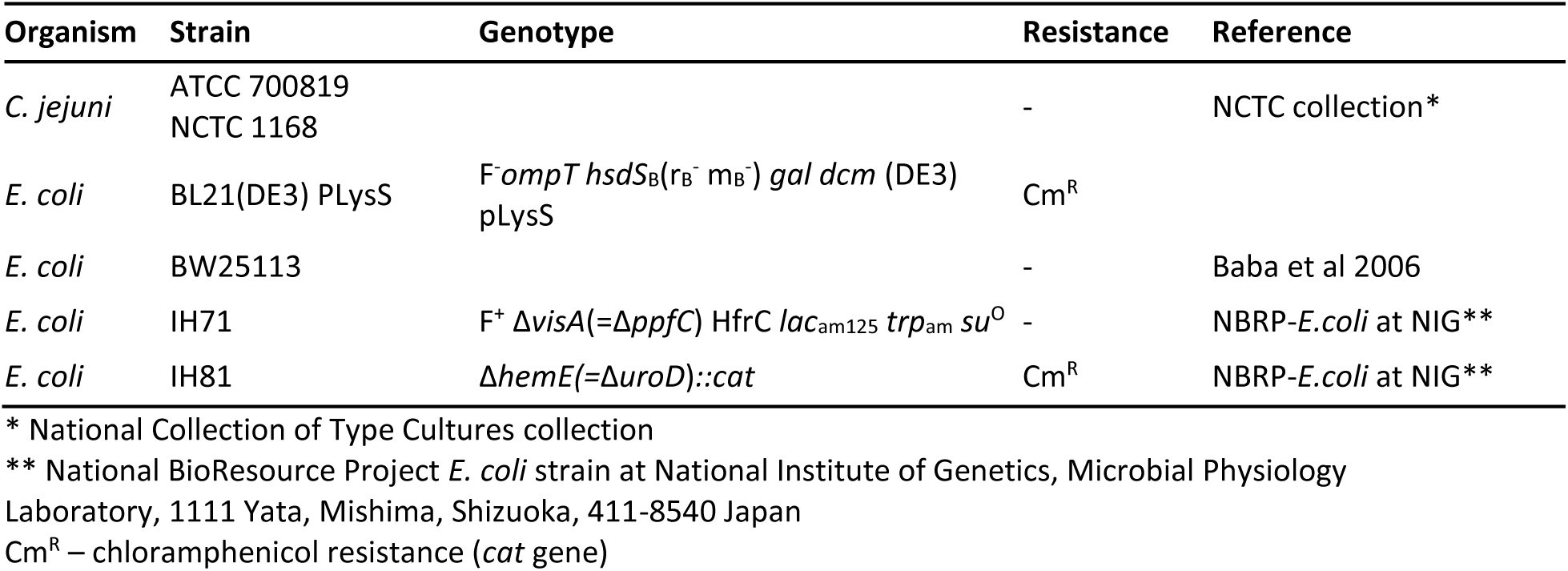
Strains used in this study

**Table S2.**
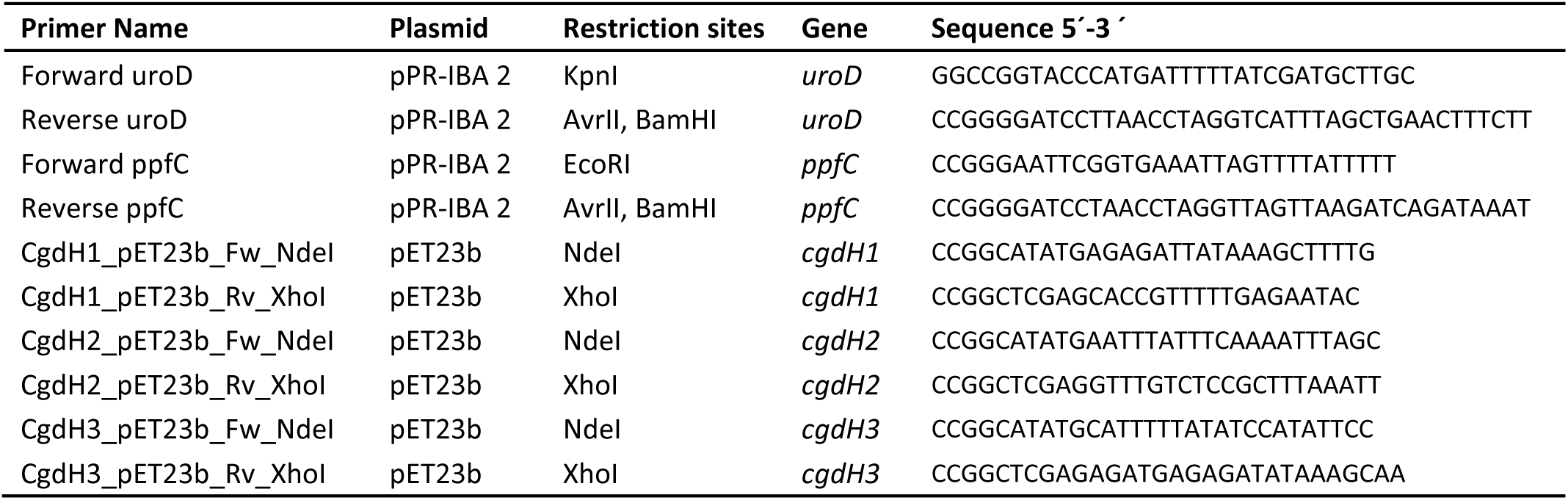
Primers used in this study

**Table S3.**
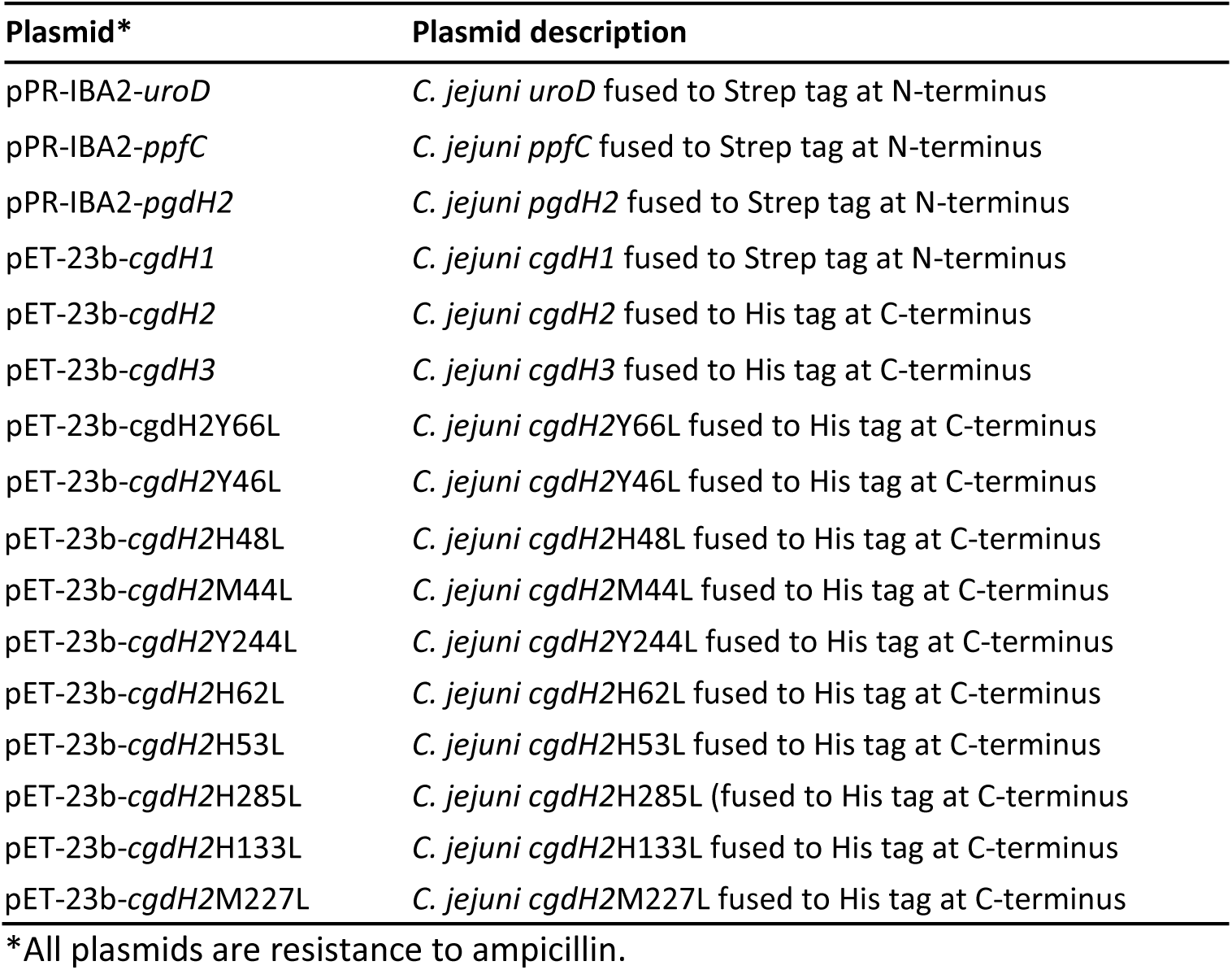
Plasmids used in this study

| Plasmid* | Plasmid description |
| --- | --- |
| pPR-IBA2-*uroD* | *C. jejuni uroD* fused to Strep tag at N-terminus |
| pPR-IBA2-*ppfC* | *C. jejuni ppfC* fused to Strep tag at N-terminus |
| pPR-IBA2-*pgdH2* | *C. jejuni pgdH2* fused to Strep tag at N-terminus |
| pET-23b-*cgdH1* | *C. jejuni cgdH1* fused to Strep tag at N-terminus |
| pET-23b-*cgdH2* | *C. jejuni cgdH2* fused to His tag at C-terminus |
| pET-23b-*cgdH3* | *C. jejuni cgdH3* fused to His tag at C-terminus |
| pET-23b-*cgdH2Y66L* | *C. jejuni cgdH2Y66L* fused to His tag at C-terminus |
| pET-23b-*cgdH2Y46L* | *C. jejuni cgdH2Y46L* fused to His tag at C-terminus |
| pET-23b-*cgdH2H48L* | *C. jejuni cgdH2H48L* fused to His tag at C-terminus |
| pET-23b-*cgdH2M44L* | *C. jejuni cgdH2M44L* fused to His tag at C-terminus |
| pET-23b-*cgdH2Y244L* | *C. jejuni cgdH2Y244L* fused to His tag at C-terminus |
| pET-23b-*cgdH2H62L* | *C. jejuni cgdH2H62L* fused to His tag at C-terminus |
| pET-23b-*cgdH2H53L* | *C. jejuni cgdH2H53L* fused to His tag at C-terminus |
| pET-23b-*cgdH2H285L* | *C. jejuni cgdH2H285L* (fused to His tag at C-terminus |
| pET-23b-*cgdH2H133L* | *C. jejuni cgdH2H133L* fused to His tag at C-terminus |
| pET-23b-*cgdH2M227L* | *C. jejuni cgdH2M227L* fused to His tag at C-terminus |

\*All plasmids are resistance to ampicillin.

**Table S4.**
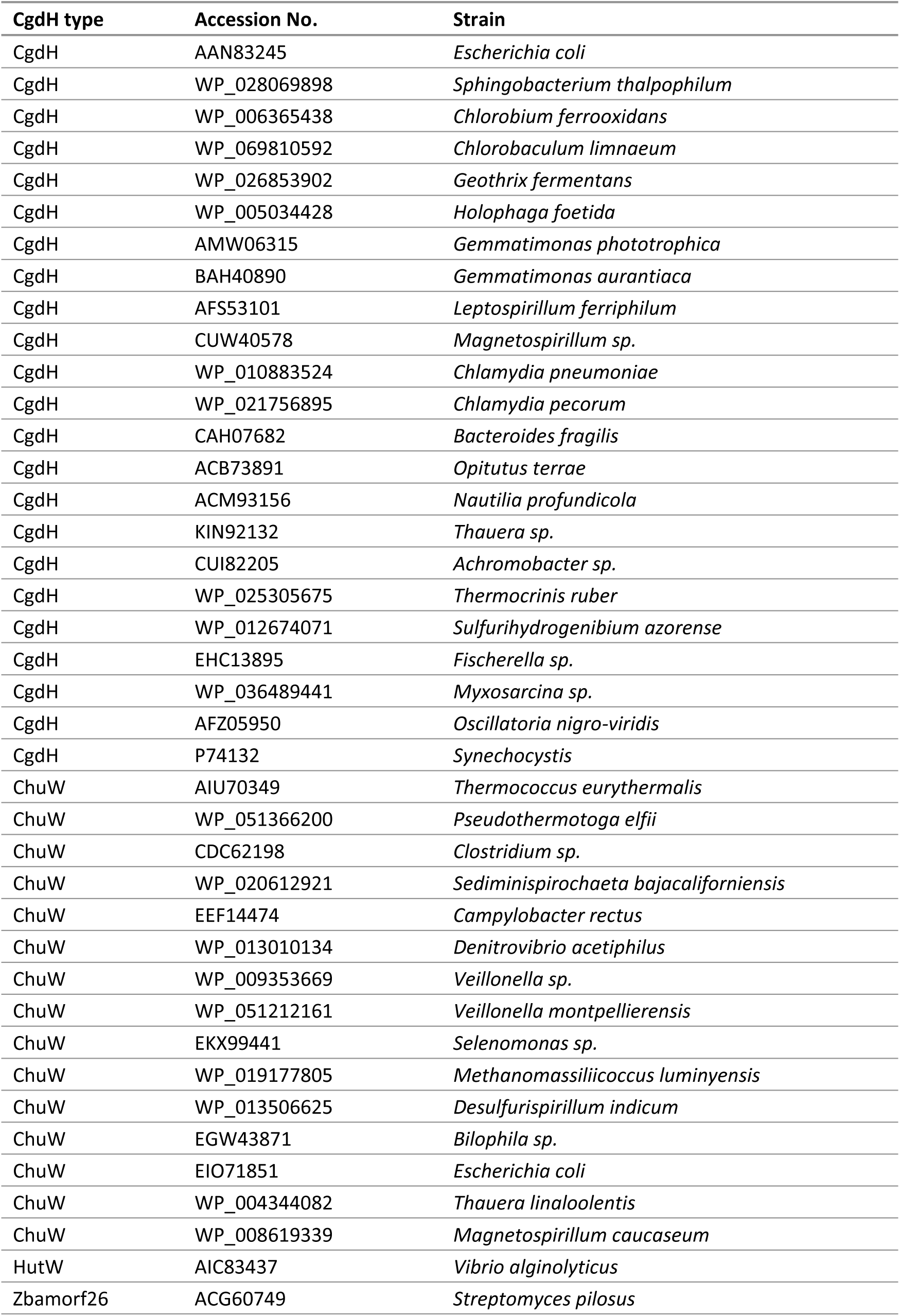
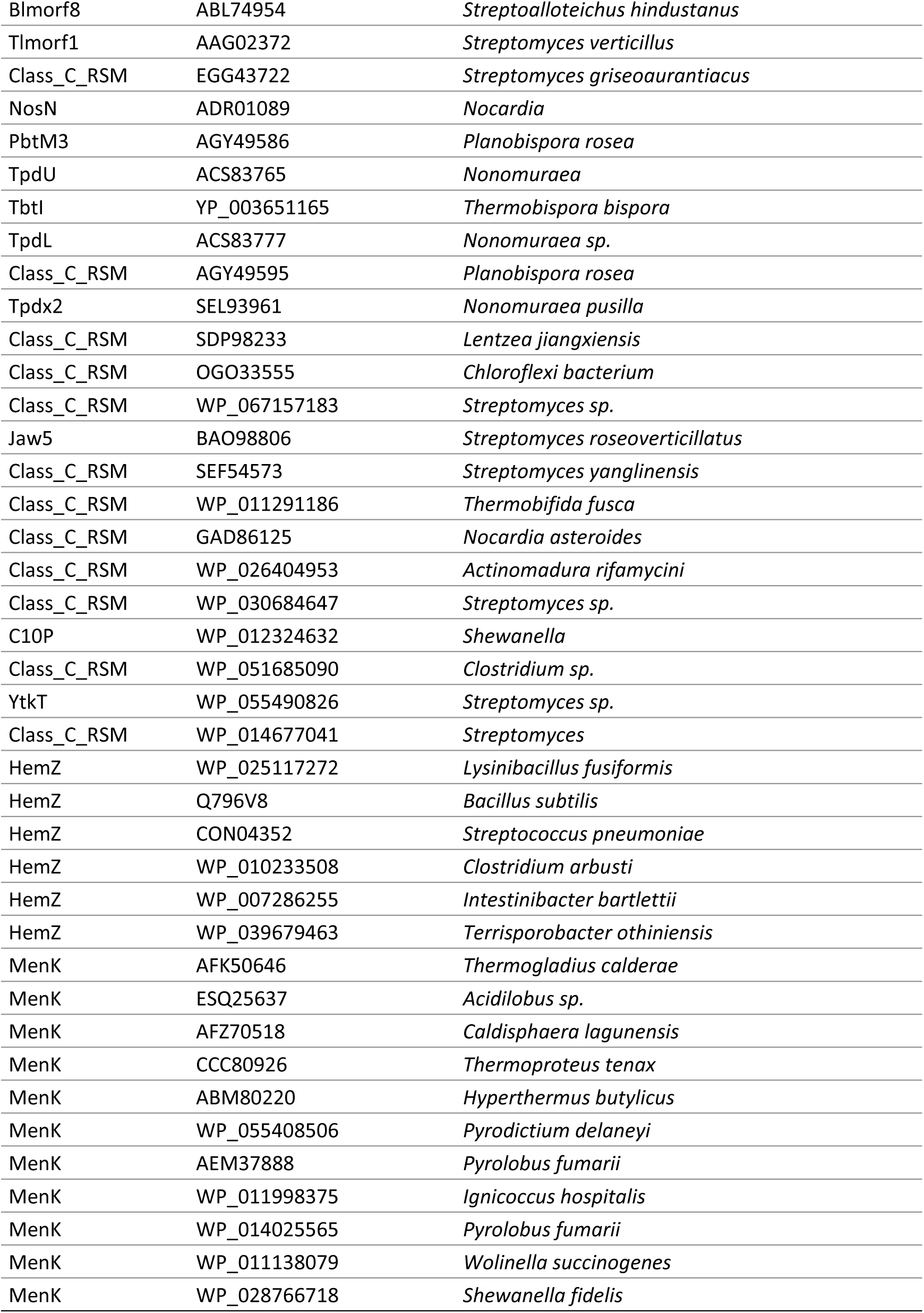
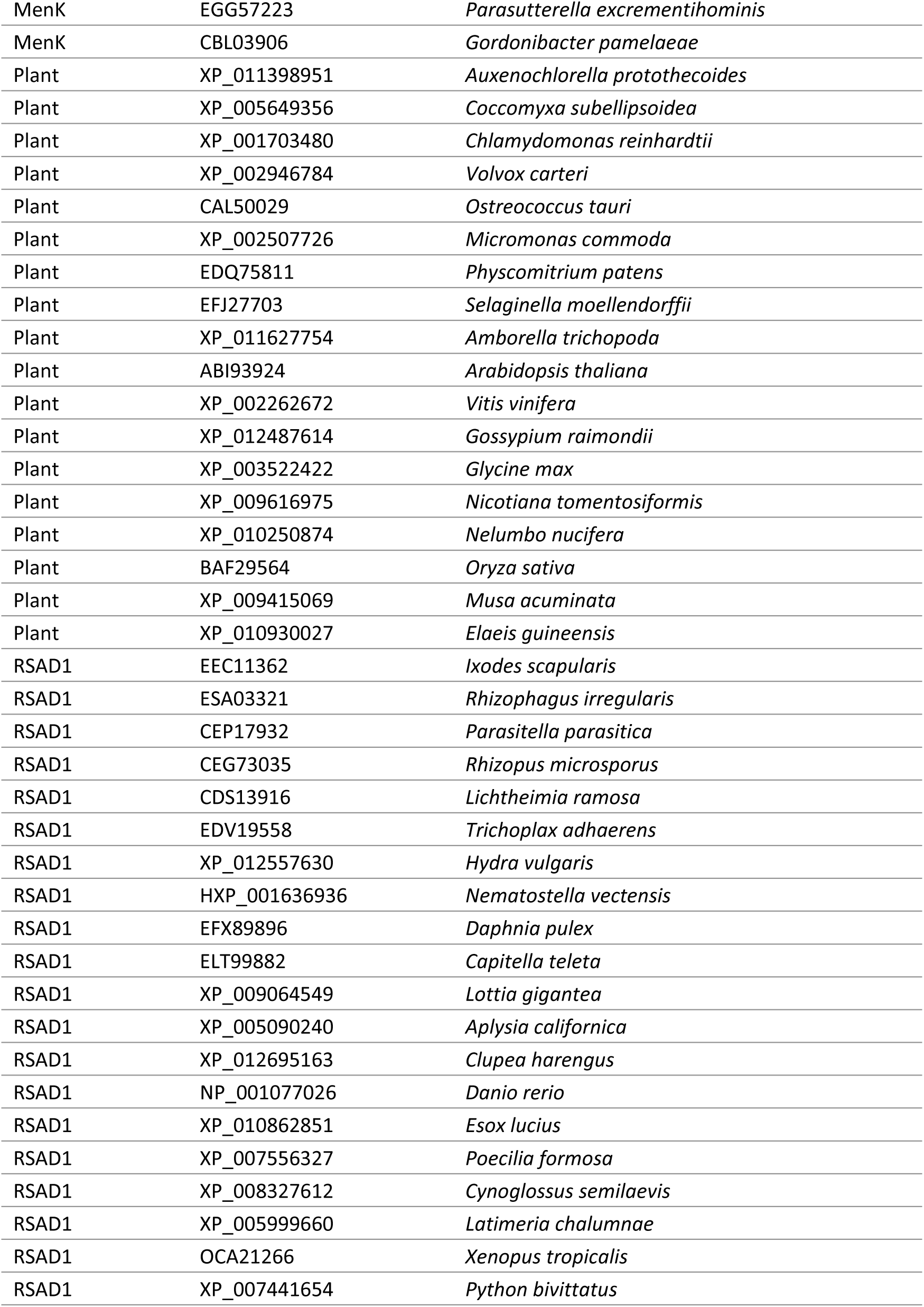
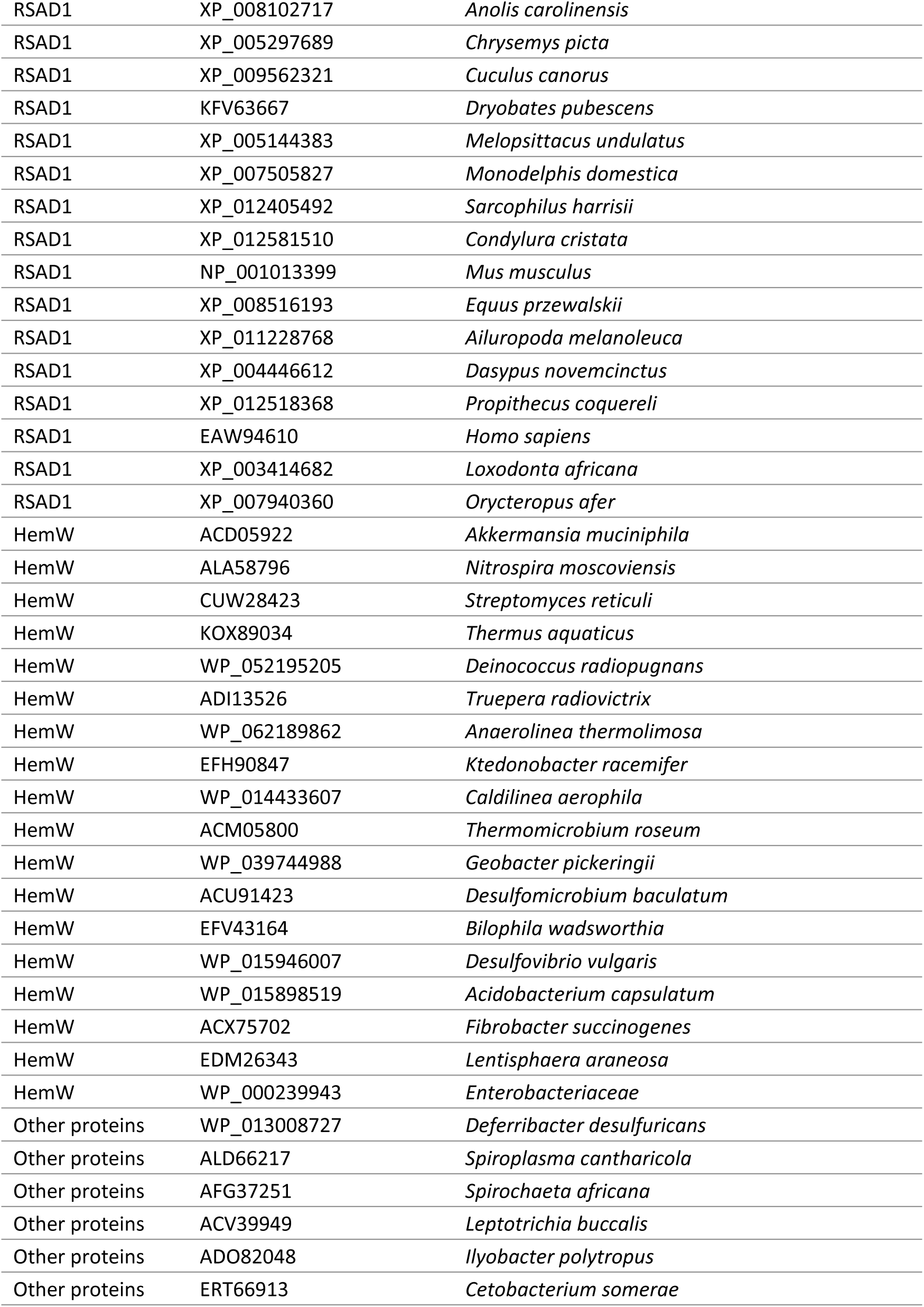
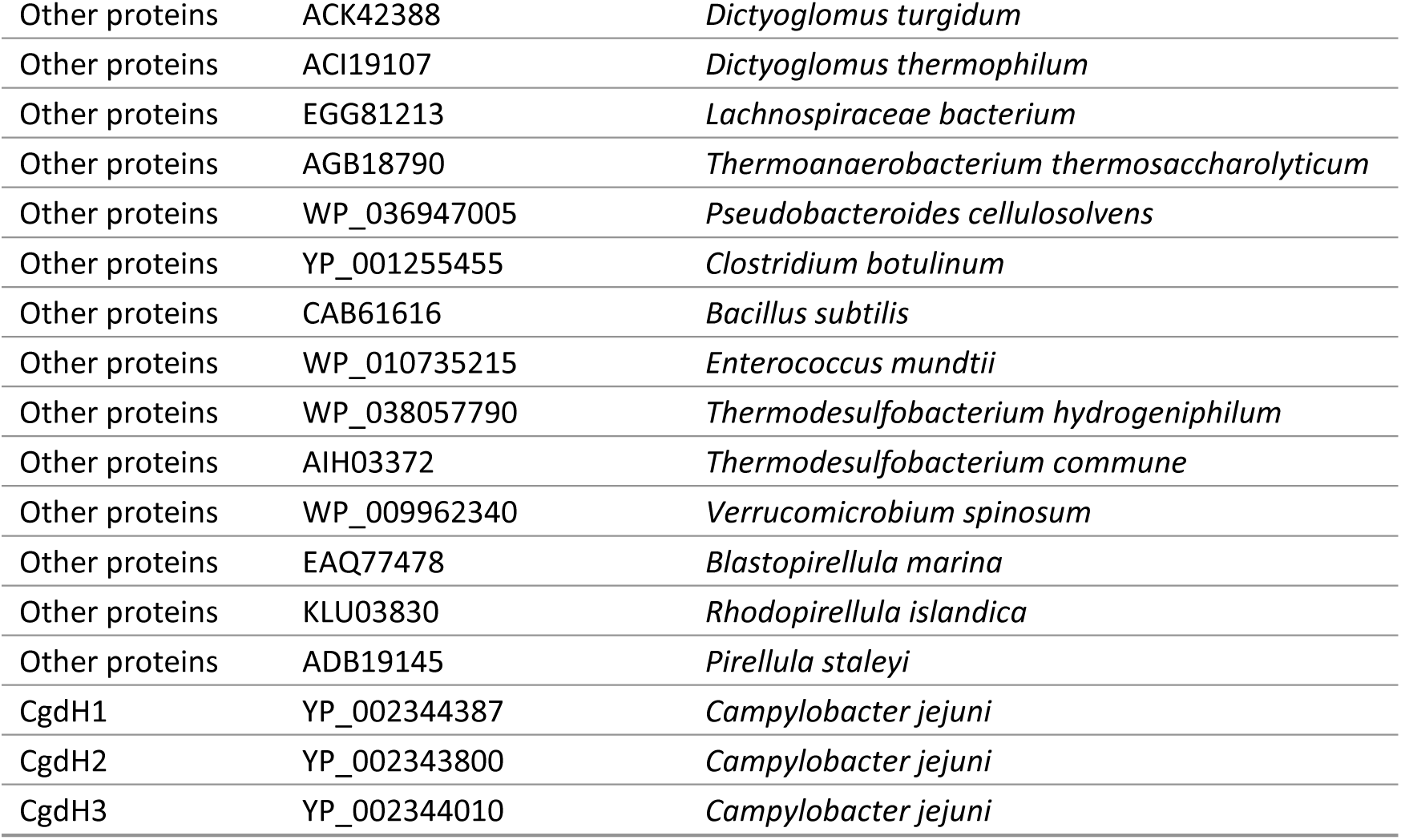
List of proteins used for the phylogenetic tree

